# Eukaryotic Elongation Factor 2 Kinase EFK-1/eEF2K promotes starvation resistance by preventing oxidative damage in *C. elegans*

**DOI:** 10.1101/2024.03.20.585993

**Authors:** Junran Yan, Forum Bhanshali, Chiaki Shuzenji, Tsultrim T. Mendenhall, Xuanjin Cheng, Pamela Bai, Gahan Diwan, Donna Seraj, Joel N. Meyer, Poul H. Sorensen, Jessica H. Hartman, Stefan Taubert

## Abstract

Cells and organisms frequently experience starvation. To adapt and survive, they mount an evolutionarily conserved stress response. A vital component in the mammalian starvation response is eukaryotic elongation factor 2 (eEF2) kinase (eEF2K), which responds to starvation by phosphorylating and inactivating the translation elongation driver eEF2, thus shutting down translation and facilitating survival. *C. elegans efk-1/eEF2K* phosphorylates EEF-2/eEF2 on a conserved residue and is required for starvation survival, but how it promotes survival remains unclear. Surprisingly, we found that eEF2 phosphorylation is unchanged in starved *C. elegans*, suggesting that *efk-1* promotes survival via a noncanonical pathway. We show that *efk-1* upregulates transcription of the DNA repair pathways, nucleotide excision repair (NER) and base excision repair (BER), to promote starvation survival. Furthermore, *efk-1* suppresses oxygen consumption and ROS production in starvation to prevent oxidative stress. Thus, *efk-1* enables starvation survival by protecting animals from starvation-induced oxidative damage through a translation-independent pathway.

## Introduction

One of the most common stresses in nature is starvation. The ability to adapt to starvation is essential for the survival of cells, tissues, and organisms. When challenged by starvation, organisms mount an evolutionarily conserved molecular and physiological response that involves the rewiring of gene expression and metabolism. In free-living animals such as the nematode worm *Caenorhabditis elegans*, timely activation of the starvation response promotes stress survival and increases evolutionary fitness. Similarly, in poorly vascularized tumors, cancer cells hijack the conserved starvation response to survive and proliferate in nutrient-poor conditions. Therefore, understanding the mechanism and regulation of the starvation response has fundamental biological and biomedical relevance.

A major mechanism of starvation adaptation involves attenuating the energy-demanding process of mRNA translation (1,2). Protein synthesis represents 30∼35% of the cellular ATP economy, of which >99% is used for translation elongation (3). The rate of translation elongation is regulated by eukaryotic elongation factor 2 kinase (eEF2K), which is also a major regulator of starvation survival (4,5). In starved mammalian cells, eEF2K attenuates protein synthesis by inactivating eukaryotic elongation factor 2 (eEF2), the rate-limiting driver of translation elongation. Specifically, eEF2K inhibits eEF2 by phosphorylating a conserved Thr56, which disrupts eEF2’s normal association with the ribosome complex (6,7). eEF2K activation thus allows starved cells to conserve energy for survival (5). eEF2K is aberrantly upregulated and promotes survival in nutrient-poor solid tumors, leading to poor patient prognoses (2,5,8–10). Conversely, eEF2K inhibition blocks cancer cell migration and tumor growth (8,11–13). However, it is unclear how eEF2K regulates downstream cytoprotective processes to promote starvation survival.

In addition to its role in translation regulation, evidence has recently emerged that eEF2K may act via alternative mechanisms. In cancer cells, eEF2K directly interacts with and phosphorylates the metabolic regulator Pyruvate kinase isozymes M2 (PKM2) in a translation-independent manner (14). Mammalian eEF2K also phosphorylates several other proteins (15), suggesting a broader regulatory role, although these interactions and their relevance have not yet been explored in detail.

To uncover regulatory roles of EFK-1/eEF2K in stress response, we used the animal model *C. elegans*, where eEF2K is evolutionarily conserved and plays important roles in stress adaptation. The *C. elegans* eEF2K ortholog EFK-1 shares sequence homology in its catalytic domain, and phosphorylates the *C. elegans* eEF2 ortholog, EEF-2, on the equivalent Thr56 residue (16,17). Similar to eEF2K^-/-^ mammalian cells (5), *C. elegans efk-1(ok3609)* loss-of-function mutants completely lack EEF-2 Thr56 phosphorylation (16,17) and exhibit impaired starvation survival (5). This suggests that *C. elegans* EFK-1 acts via EEF-2 and translation modulation to promote organism survival in starvation, similar to the mechanism described in mammalian cells.

To study how *efk-1* promotes starvation survival, we used the *efk-1(ok3609)* mutant which we confirmed was functionally null and had diminished *efk-1* mRNA expression, abolished EEF-2 Thr56 phosphorylation, and defective starvation survival. Surprisingly, unlike in mammalian cells, EEF-2 phosphorylation is constitutively high in both fed and starved wild-type *C. elegans*, suggesting an alternative, translation-independent mechanism of *efk-1*. To map this noncanonical pathway, we identified two transcription factors, bZIP transcription factor family 2 (ZIP-2) and *C. elegans* p53-like protein 1 (CEP-1), which are also required for starvation resistance and function in the *efk-1* pathway. Transcriptomic profiling of *efk-1*, *zip-2*, and *cep-1* mutants revealed that the three factors are jointly required for increased expression of DNA repair pathways during starvation. Specifically, *efk-1* is required to upregulate nucleotide excision repair (NER) and base excision repair (BER) to increase resistance to oxidative DNA damage, which is linked to starvation-induced oxidative stress. Additionally, *efk-1* prevents reactive oxygen species (ROS) accumulation in the cell and represses oxygen consumption, all of which may ameliorate oxidative stress during starvation. As these cytoprotective effects of *efk-1* are independent from translation elongation regulation and phosphorylation status of EEF-2, our studies reveal a new, noncanonical mechanism of *efk-1*-mediated starvation resistance.

## Results

### *efk-1(ok3609)* is a null allele of *efk-1*

The *efk-1(ok3609)* mutant shows deficiency in starvation survival (5), but its molecular lesion has not been precisely defined. Sequencing the *efk-1(ok3609)* loss-of-function allele revealed that *ok3609* is an in-frame deletion spanning the 3’ half of exon 2 and the beginning of exon 3, resulting in the loss of a part of the conserved alpha-kinase domain of EFK-1 (Figure 1A). In line with loss of kinase activity, phosphorylation of Thr56 of EEF-2, the evolutionarily conserved substrate of EFK-1, is abolished in *efk-1* mutants (Figure 1B; Figure S1A-B), as previously shown (16,17). Thus, like mammalian eEF2K (5,7), *C. elegans efk-1* appears to be the sole kinase that phosphorylates Thr56 of EEF-2. Although the deletion in *efk-1(ok3609)* is in frame for both *efk-1* isoforms (Figure 1A), RT-qPCR revealed that *efk-1* mutants have approximately 10-fold decreased *efk-1* transcript expression (Figure 1C). We conclude that *ok3609* is a null allele of *efk-1*.

**Figure 1.**
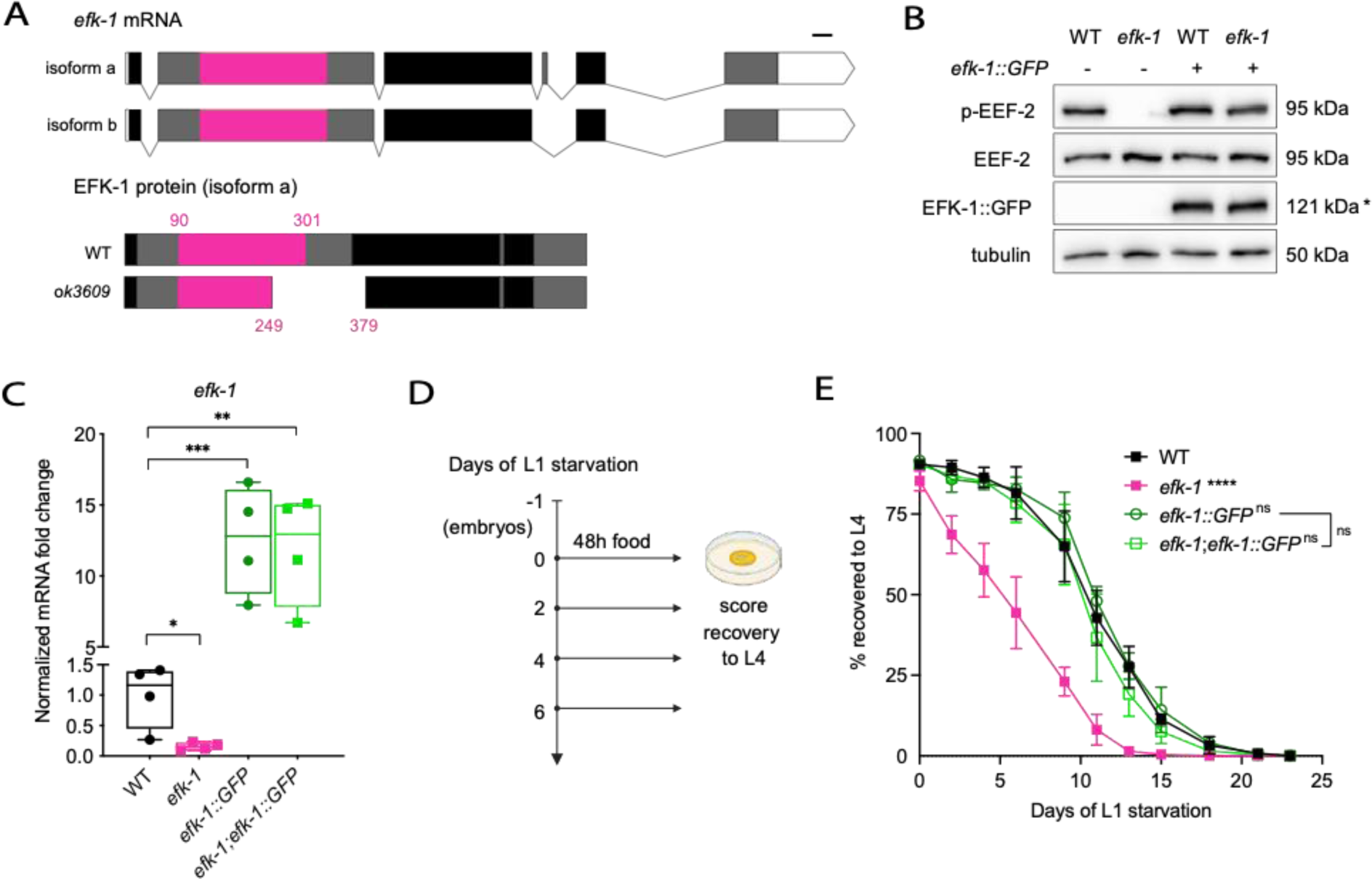
*ok3609* is a null allele of *efk-1*. **(A)** Sequence map of *efk-1* mRNA (top) and EFK-1 protein (bottom). Red, alpha-kinase domain; white, UTR; exons, black and grey. Scale bar, 100 bases or 33 amino acids. **(B)** Western Blot (WB) of EEF-2 Thr56 phosphorylation (p-EEF-2), EEF-2, GFP and tubulin in wild type (WT) and *efk-1(ok3609)* mutants with or without the rescue construct *efk-1::GFP*. Asterisk (*) denotes predicted protein size. N=3; p-EEF-2 is quantified in Figure S1A; for full membrane see Figure S1B. **(C)** Real-time quantitative PCR (RT-qPCR) of *efk-1* mRNA expression in WT and *efk-1* mutants with or without the rescue construct *efk-1::GFP*. N=4; *p<0.05, ** p<0.01, ***p<0.001 (unpaired t-test). **(D)** Schematic of L1 starvation survival experiment. **(E)** The graph shows population viability (percent able to reach at least the L4 stage, y-axis) over time (days of L1 stage starvation, x-axis) of WT and *efk-1(ok3609)* mutants with or without the rescue construct *efk-1::GFP*. N=4, error bars represent standard deviation (SD); ****p<0.0001 percent L4 vs. WT animals (AUC compared using one-way ANOVA with Tukey’s multiple comparisons test). WT, wild-type; ns, not significant; AUC, area under the curve. See Source data for **(C, E)**.

*efk-1* is required for survival and recovery from starvation-induced arrest at the first larval stage (5) (Figure 1D-E). To ascertain that the starvation sensitivity of the *efk-1(ok3609)* mutant is due to loss of *efk-1* function and to study the expression of EFK-1 *in vivo*, we generated a strain that stably expresses a translational *efk-1::GFP* fusion protein from its own promoter (*efk-1p::EFK-1::GFP*; see Methods). In L4 stage worms, *efk-1::GFP* is expressed mainly in the cytoplasm of the hypodermis, neurons, and intestinal cells (Figure S1C). In the *efk-1(ok3609)* mutant background, this transgene restored both EEF-2 Thr56 phosphorylation and L1 starvation survival to wild-type levels (Figure 1B, E), indicating complete functional rescue. RT-qPCR analysis revealed approximately 10-fold higher *efk-1* mRNA expression in this strain compared to wild type (Figure 1C). However, compared to wild type, the *efk-1::GFP* strain did not show increased substrate phosphorylation (Figure 1B; Figure S1A), nor did it show increased starvation resistance (Figure 1E). These data suggest that the *efk-1::GFP* strain does not have gain-of-function activity. In sum, we validated that *efk-1(ok3609)* is a null allele of *efk-1*, and that *efk-1* is required for starvation survival and EEF-2 T56 phosphorylation.

### *efk-1* promotes starvation resistance via a translation-independent mechanism

In mammalian cells, eEF2K phosphorylates and inactivates EEF2 in nutrient deprivation and hypoxia, which attenuates translation elongation, reduces energy expenditure, and promotes cell survival (5,18). In *C. elegans*, starvation also leads to global translation attenuation (19), suggesting that EFK-1–EEF-2 might function in this nematode as it does in mammals. To test if *C. elegans efk-1* attenuates translation elongation in starvation, we examined EEF-2 phosphorylation (p-EEF-2) in starved L1 and L4 larvae with Western blots. As expected (16,17), EEF-2 Thr56 phosphorylation was abolished in *efk-1* mutants, both in L4 (Figure 2A; Figure S2A-C) and L1 larvae (Figure 2B; Figure S2D-F) stages. In fed *efk-1* mutants, we also observed increased basal levels of eIF2α phosphorylation (p-eIF2α), a regulatory mark of translation initiation, possibly indicating compensation for the lack of EEF-2 phosphorylation (Figure S2B, E). Surprisingly, however, we did not observe an increase in p-EEF-2 levels in wild-type L4 worms subjected to eight hours of starvation (Figure 2A; Figure S2A) or in L1 worms subjected to prolonged starvation of up to nine days (Figure 2B; Figure S2D). Instead, p-EEF-2 levels were already substantial at baseline in well-fed L1 and L4 worms (Figure 2C; Figure S2G-H), with no apparent increase in starvation. In fact, p-EEF-2 levels decreased with starvation (Figure S2A, D). This high baseline level of p-EEF-2 is consistent with previous Western blot and phosphoproteomics studies in *C. elegans* (16,17,20). In contrast, in A549 lung cancer cells, we observed low or absent p-eEF2 signals in fed conditions and substantial induction after 3 to 6 hours of starvation (Figure 2D; Figure S2I-J), consistent with studies in multiple mammalian cell lines (5,21). Thus, although canonical EFK-1–EEF-2 signaling is clearly present in *C. elegans* (Figure 2A-B), *efk-1* surprisingly does not appear to mediate starvation resistance by increasing EEF-2 phosphorylation.

**Figure 2.**
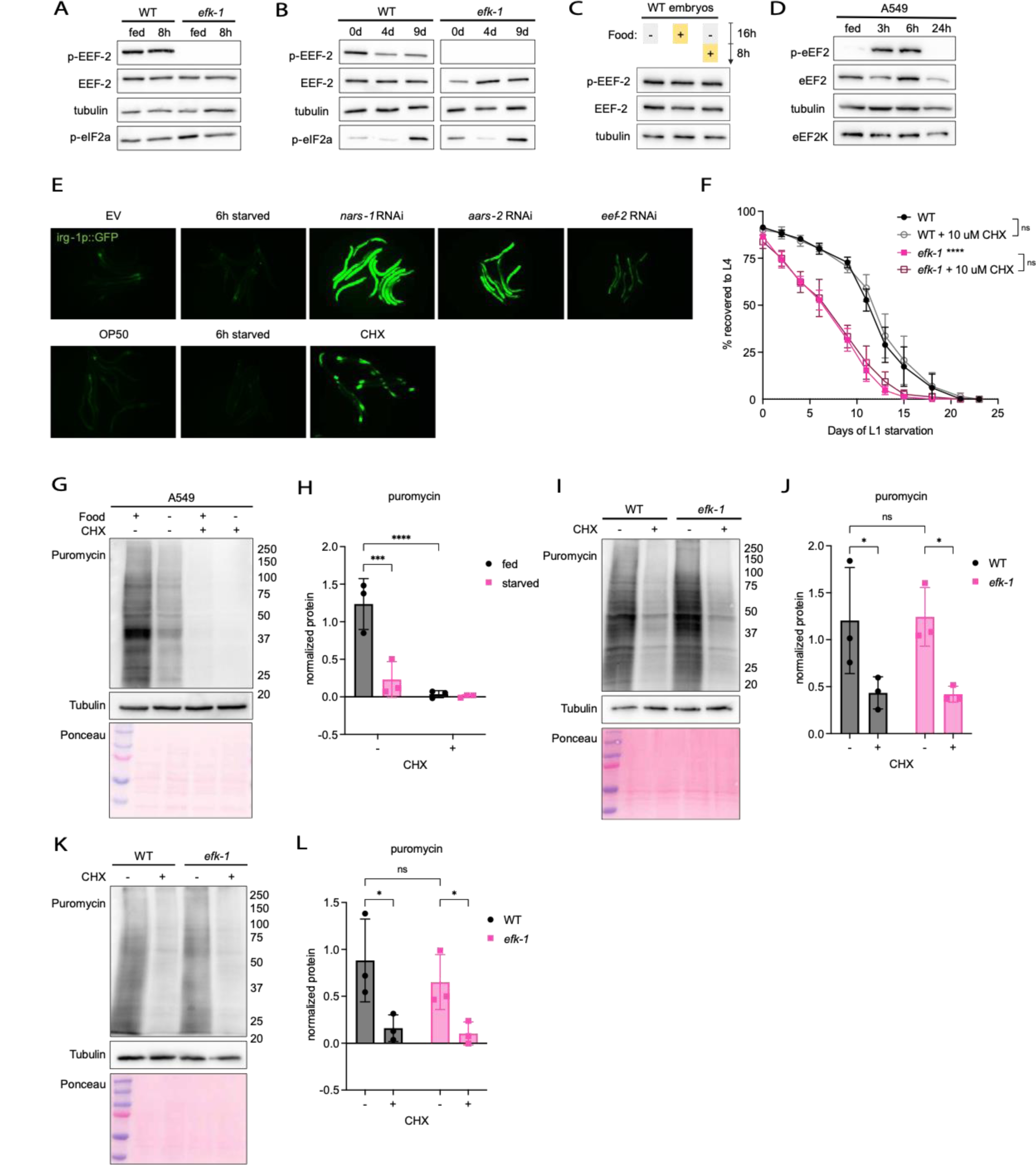
*efk-1* promotes starvation survival via a noncanonical mechanism. **(A-B)** WB of p-EEF-2, EEF-2, tubulin and eIF2α Ser51 phosphorylation (p-eIF2α) in WT and *efk-1* mutants under the following conditions: **(A)** fed and 8-hour starved in L4 stage larvae, and **(B)** overnight hatched (0d), 4-days starved (4d) and 9-days starved (9d) at L1 stage. N=4; p-EEF-2 and p-eIF2α quantified in Figure S2A-B, D-E; for full membrane see Figure S2C, F. **(C)** WB of p-EEF-2, EEF-2 and tubulin in overnight hatched and various fed conditions in L1-stage WT worms. Worms were either hatched without food, hatched in food, or hatched without food and refed for 8 hours. N=3; p-EEF-2 quantified in Figure S2G; for full membrane see Figure S2H. **(D)** WB of eEF2 Thr56 phosphorylation (p-eEF2), eEF2, tubulin, and eEF2K in A549 lung cancer cells in fed condition and after serum/glucose deprivation for 3, 6, and 24 hours. N=4; p-eEF2 quantified in Figure S2I; for full membrane see Figure S2J. **(E)** Fluorescence micrographs of *irg-1p::GFP* reporter worms (L4 stage) after RNAi against various translation machinery components or treatment with 2 mg/ml cycloheximide for 3h. For RNAi experiments, empty vector RNAi-expressing HT115 *E. coli* (EV) is used as control. Data shown are representative of three independent experiments. **(F)** The graph shows L1 starvation survival of WT and *efk-1* mutants with or without supplementation of 10 uM cycloheximide (CHX). N=4, error bars represent SD; ****p<0.0001 percent L4 vs. WT animals (AUC compared using one-way ANOVA with Tukey’s multiple comparisons test). **(G-H)** WB and quantification of puromycin incorporation in A549 lung cancer cells. Cells were treated with 1 ug/ml puromycin for 10 minutes. 1 hour of 2 ug/ml CHX is used as control. N=3, error bars represent SD; ***p<0.001, ****p<0.0001 (two-way ANOVA with uncorrected Fisher’s LSD); for full membrane see Figure S2M. **(I-L)** WB and quantification of puromycin incorporation in *efk-1* mutants and WT both in **(I-J)** 8-hour L4 starvation, and **(K-L)** 6-day L1 starvation. Worms were treated with 0.5 mg/ml puromycin for 3 hours. 6 hours of 2 mg/ml CHX is used as control. N=3, error bars represent SD; *p<0.05 (two-way ANOVA with uncorrected Fisher’s LSD); for full membrane see Figure S2N-O. WT, wild-type; ns, not significant; AUC, area under the curve. See Source data for **(F, H, J, L)**.

To further study whether starvation arrests translation elongation in a canonical, EEF-2 dependent manner, we assessed a downstream readout of translation elongation arrest. Conditions of translation elongation arrest, including RNA interference (RNAi) knockdown of EEF-2 or amino-acyl tRNA synthetases, activates transcription of a downstream gene, *irg-1* (22,23). As EFK-1 is solely responsible for EEF-2 Thr56 phosphorylation and is a potent inhibitor of EEF-2 activity (Figure 2A-B), *irg-1* expression should be activated if starvation were to arrest translation elongation via the EFK-1–EEF-2 axis. To test this, we starved worms expressing a transcriptional *irg-1p::GFP* promoter reporter (24). In line with the lack of increased EEF-2 phosphorylation in starvation (Figure 2A-B), we did not observe increased *irg-1* promoter activity during starvation (Figure 2E), consistent with published RNAseq studies (20). In contrast, translation arrest via cycloheximide (CHX) treatment (25), *eef-2* RNAi, or RNAi against amino-acyl tRNA synthetases *nars-1* and *aars-2* strongly induced *irg-1* promoter activity (Figure 2E), as reported (22,26). The lack of *irg-1* induction in starvation is consistent with the above data that EFK-1 does not appear to increase EEF-2 phosphorylation in starvation, supporting the hypothesis that EFK-1 functions in starvation in a noncanonical pathway.

Next, we asked whether *efk-1* promotes starvation survival via inhibiting translation elongation, and whether *efk-1* loss causes starvation sensitivity by derepressing translation elongation. In cancer cells, the defective starvation survival of eEF2K-inactive cells is rescued by the pharmacological translation elongation inhibitor CHX, demonstrating that eEF2K promotes starvation survival mainly via translation elongation inhibition (21). Using the same approach in *C. elegans*, we investigated whether CHX is sufficient to rescue the starvation survival defect of *efk-1* mutants. However, we found that various doses of CHX did not improve the starvation survival defect of *efk-1* mutants (Figure 2F; Figure S2K-L). This was unlikely due to poor CHX uptake or activity, as similar doses of CHX extend worm lifespan (27), and higher doses substantially reduce translation and induce the *irg-1p::gfp* transcriptional reporter (Figure 2E), as published (22,26,28). The fact that CHX supplementation does not rescue the starvation survival defect of *efk-1* mutants suggests that their starvation sensitivity is not due to derepressed translation, consistent with the above findings that *efk-1* is unlikely to promote starvation survival by inhibiting translation elongation.

Finally, to directly assess if *efk-1* inhibits global protein synthesis in starving worms, we assayed global protein synthesis rates using the surface sensing of translation (SUnSET) assay. SUnSET measures *de novo* protein synthesis rates by puromycin labeling of newly synthesized peptides (29–31), and has been widely used in mammalian cells and *C. elegans* (26,32,33,28). In cell lines, puromycin incorporation is decreased in starvation and other conditions with active eEF2K (34,35). Indeed, we detected decreased protein synthesis in A549 cells after six hours of starvation (Figure 2G-H; Figure S2M), as previously shown (34). Next, we used SUnSET to test whether *C. elegans efk-1* mutants derepress translation when subjected to 8-hour starvation at L4 and 6-day starvation at L1. Interestingly, we did not detect increased total puromycin incorporation in *efk-1* mutants compared to wild type in either condition (Figure 2I-L; Figure S2N-O). This suggests that *efk-1* does not suppress global translation during starvation, consistent with the above data that p-EEF-2 remained unchanged in the same conditions. Taken together, these data suggest that EFK-1 promotes starvation resistance via a noncanonical pathway independent from EEF-2 inhibition and global translation attenuation.

### Transcription factors ZIP-2/bZIP and CEP-1/p53 function in the *efk-1* starvation response pathway

Because *efk-1* regulates starvation survival in a translation-independent manner, we hypothesized that it may act via transcriptional regulation. Starvation arrest induces extensive transcriptomic rewiring to repress many genes while activating specific genes required for mounting the starvation response (36). Accordingly, starvation survival in *C. elegans* requires numerous transcription factors (TFs), including insulin signaling component *daf-16/ Forkhead box O (FOXO)*, and metabolic regulators *hlh-30/ Transcription factor EB (TFEB)*, nuclear hormone receptor 49 (*nhr-49*), *zip-2/bZIP*, and *cep-1/p53* (36–42). To delineate how *efk-1* regulates downstream gene expression, we searched for starvation-responsive transcription factors with functional links to *efk-1* or mammalian eEF2K. Interestingly, EEF-2 is inactivated not only during starvation by EFK-1, but also after pathogen exposure by bacterial endotoxins. This inactivation triggers a host immune response that is orchestrated by the TFs ZIP-2/bZIP and CEBP-2/ CCAAT Enhancer Binding Protein Gamma CEBPG (22,24,43–45). *zip-2* is also required for starvation survival and dietary restriction-induced longevity (39,46). In addition, the *C. elegans* p53 ortholog *cep-1* is required for starvation survival (47,48,41), and human p53 interacts genetically with eEF2K in tumor cells (49). Thus, we hypothesized that *zip-2*, *cebp-2*, and/or *cep-1* might function with *efk-1* to promote starvation survival.

To study the roles of these TFs in the starvation response, we quantified the population survival of previously characterized *zip-2*, *cep-1*, and *cebp-2* loss-of-function mutants (22,41,43,50,51) by L1 starvation survival assays. As expected, *zip-2*, *cebp-2*, or *cep-1* loss caused starvation sensitivity and thus phenocopies *efk-1* loss (Figure 3A) (5). *cebp-2* mutants also showed growth defects under standard growth conditions, whereas *zip-2* and *cep-1* mutants did not (Figure S3A). To examine the genetic interactions of *efk-1* with *zip-2*, *cebp-2*, and *cep-1*, we next constructed respective double mutants. In standard growth conditions, *zip-2;efk-1* and *cep-1;efk-1* mutants were viable without apparent phenotypes (Figure S3B), whereas *efk-1*;*cebp-2* mutants could not be obtained, suggesting synthetic lethality/sterility. Interestingly, *zip-2;efk-1* and *cep-1;efk-1* double mutants showed no additive defect in L1 starvation survival compared to respective single mutants (Figure 3B-C). Because *zip-2* and *cep-1* both genetically interact with *efk-1*, we next constructed a *zip-2;cep-1* double mutant, which also showed no additive starvation survival defect compared to the respective single mutants (Figure 3D; Figure S3C). Thus, *zip-2* and *cep-1*, but not *cebp-2*, are components of the *efk-1* starvation response pathway.

**Figure 3.**
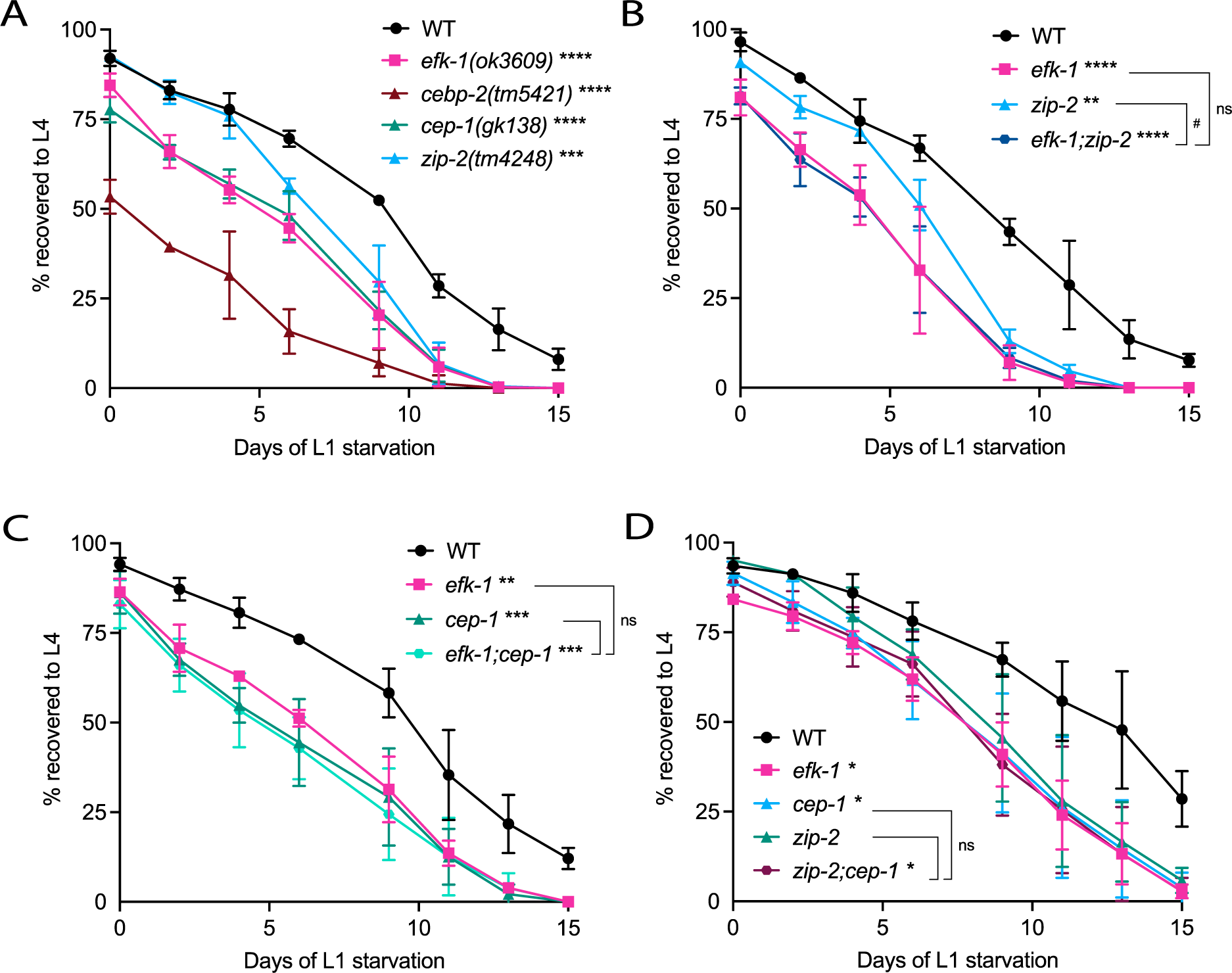
The TFs *zip-2* and *cep-1* are required in the *efk-1* pathway for starvation survival. **(A)** The graph shows L1 starvation survival of WT, *efk-1*, *zip-2(tm4248)*, *cebp-2(tm5421)*, and *cep-1(gk138)* single mutants. N=3, error bars represent SD; ***p<0.001, ****p<0.0001 percent L4 vs. WT animals (AUC compared using one-way ANOVA with Dunnett’s multiple comparisons test). **(B-D)** The graphs show the L1 starvation survival of **(B)** *efk-1;zip-2*, **(C)** *efk-1;cep-1* and **(D)** *zip-2;cep-1* double mutants alongside the respective single mutants. N=3-4, error bars represent SD; *p<0.05, **p<0.01, ***p<0.001, ****p<0.0001 percent L4 vs. WT animals, ^#^p<0.05 vs. *zip-2* animals (AUC compared using one-way ANOVA with Tukey’s multiple comparisons test). WT, wild-type; ns, not significant; AUC, area under the curve. See Source data for **(A-D)**.

### DNA repair pathways are transcriptionally upregulated by *efk-1*, *zip-2*, and *cep-1* during starvation

To identify genes and pathways regulated by *efk-1*, *zip-2*, and *cep-1* in starvation, we performed RNA sequencing (RNA-seq) of fed and starved wild-type worms and *efk-1*, *zip-2*, and *cep-1* mutants (Figure 4A; Figure S4A). We found that in 8-hour starved, wild-type L4 stage worms, 3302 genes were significantly altered (p<0.005, FDR<0.05), including 1493 upregulated and 1809 downregulated genes (Table S1). To validate our expression analysis, we reanalyzed published RNA-seq data of 6-hour starved L4 stage wild-type worms (20) and compared them to our dataset. Significantly regulated genes (p<0.005 and FDR<0.05) shared a substantial overlap and correlated well between both datasets (r^2^=0.87, p<2.2e-16; Figure S4B-C). Our data also identified several experimentally confirmed starvation-induced genes, such as *fmo-2*, *cpt-3*, *fat-3*, *icl-1*, *fat-2*, *acs-11*, *lbp-1*, and *efk-1*, and starvation-repressed genes, such as *lbp-8*, *ech-6*, *acdh-1*, *acdh-2*, *acox-1.4*, *cpt-4*, and *fat-7* (5,40,52) (Figure S4D). Additionally, consistent with published transcriptomic data (20), we did not observe transcript induction of *irg-1* (Figure S4E-F), confirming that starvation does not activate *irg-1* (Figure 2E). Overall, our data recapitulate known short-term starvation responses.

**Figure 4.**
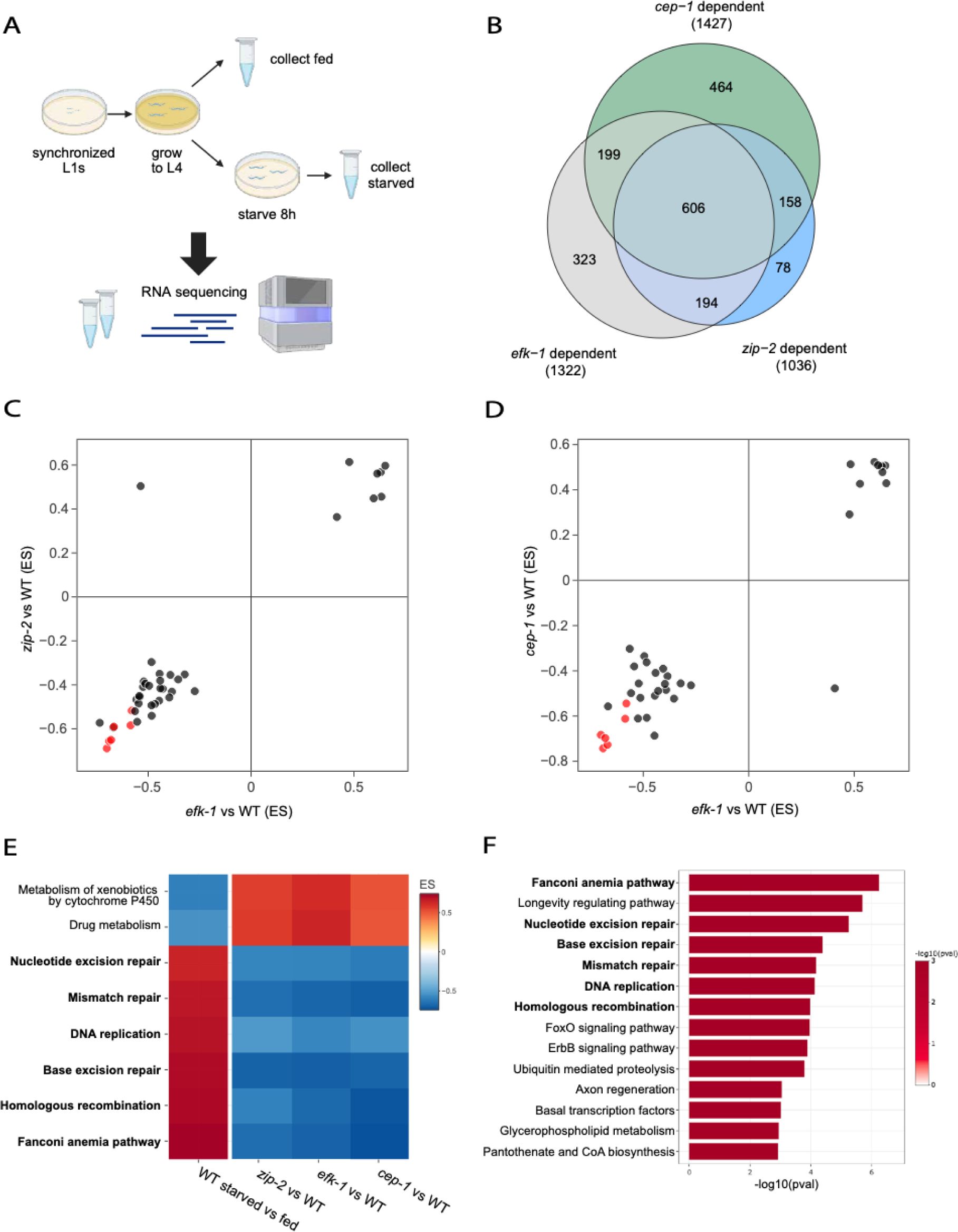
DNA repair pathways are upregulated in WT and attenuated in *efk-1*, *zip-2*, and *cep-1* mutants during starvation. **(A)** Scheme of the RNA sequencing experiment (N=3). **(B)** The Venn diagram shows the overlap between *efk-1*, *zip-2*, and *cep-1* dependent genes (p<0.005, FDR<0.05 in starved WT, but not in the respective mutant). The central intersection contains 606 *efk-1*, *zip-2*, and *cep-1* dependent genes (p<0.005, FDR<0.05 in starved WT, but not in any of *efk-1*, *zip-2*, or *cep-1* mutants). See Table S1. **(C-D)** The figure shows the correlation of KEGG pathways altered in *efk-1*, *zip-2*, and *cep-1* null mutants (pval<0.25) by GSEA, especially downregulation of DNA repair (highlighted in red). x= ES (x), y= ES (y). n=41, r^2^=0.83, p<2.2e-16 for *zip-2* vs *efk-1*; n=37, r^2^=0.87, p<2.2e-16 for *cep-1* vs *efk-1*. See Table S2. **(E)** The heatmap shows the most significantly altered pathways in *efk-1*, *zip-2*, and *cep-1* mutants relative to WT (pval<0.05, padj<0.05, |ES|>0.5 in all datasets) by GSEA. Colors represent ES. See Table S2. **(F)** The Bar plot shows functionally enriched categories in the 606 *efk-1*, *zip-2*, and *cep-1* dependent genes (pval<0.005, padj<0.0001) by ORA. Colors represent −log10(pval). DNA repair-related pathways are bolded. See Table S3. ES, enrichment score; GSEA, gene set enrichment analysis; FDR, false discovery rate; KEGG, Kyoto Encyclopedia of Genes and Genomes; ORA, overrepresentation analysis; pval, p-value; padj, adjusted p-value.

Next, we analyzed the transcriptional dysregulations in starved *efk-1*, *zip-2*, and *cep-1* mutants. We found that among the 3302 genes significantly regulated in WT, 1322 failed to be regulated in *efk-1* mutants, 1036 in *zip-2* mutants, and 1427 in *cep-1* mutants. We call these *efk-1*, *zip-2*, or *cep-1* dependent genes, respectively (Table S1). Interestingly, *efk-1*, *zip-2*, and *cep-1* mutants showed similar transcriptome deregulations in starvation compared to wild type: among the 1322 *efk-1*-dependent, 1036 *zip-2*-dependent, and 1427 *cep-1*-dependent genes, nearly half (606 genes) are dependent on all three genes (Figure 4B; Table S1). To obtain an unbiased profile of starvation gene dysregulations in these mutants, we performed second-order comparisons of *efk-1*, *zip-2*, and *cep-1* starved-versus-fed profiles to that of wild type. *efk-1* and *zip-2* mutant starvation profiles showed strong positive correlation with each other (p<0.25, r^2^=0.84, p<2.2e-16; Figure S4G), whereas *efk-1* and *cep-1* profiles exhibited a moderate positive correlation (p<0.25, r^2^=0.57, p<2.2e-16; Figure S4H). Thus, *efk-1*, *zip-2*, and *cep-1* control a shared set of genes in starved *C. elegans*.

To identify biological processes jointly regulated by *efk-1*, *zip-2*, and *cep-1* in starvation, we used two complementary approaches. First, we used gene set enrichment analysis (GSEA) to capture broad transcriptomic changes in pathways in an unbiased manner. Starving wild type worms upregulated the expression of known starvation-inducible pathways, such as “autophagy”, “FOXO signaling pathway”, and “longevity regulating pathway”, and downregulated pathways such as “drug metabolism”, “metabolism of xenobiotics by cytochrome P450”, and “oxidative phosphorylation” (Table S2), consistent with previous reports (20,36,53). Interestingly, we also observed an upregulation of DNA repair pathways, such as “nucleotide excision repair”, “base excision repair”, “homologous recombination” (HR), and “Fanconi anemia pathway” (FA) (Table S2). Next, we performed GSEA on the starvation profiles of *efk-1*, *zip-2*, and *cep-1* mutants. Similar to the gene-level correlation, the pathways enriched in *efk-1*, *zip-2*, and *cep-1* mutants correlated strongly with each other (r^2^=0.83 for *efk-1* vs. *zip-2* regulated pathways, r^2^=0.87 for *efk-1* vs. *cep-1* regulated pathways, p<2.2e-16; Figure 4C-D). Strikingly, among the pathways that were upregulated in starved wild type (p<0.05, padj<0.05, ES>0.5), DNA repair pathways such as NER, BER, HR, and FA were downregulated in all three mutants compared to wild type (p<0.05, padj<0.05, ES<-0.5) (Figure 4E).

To complement the GSEA analysis, we performed overrepresentation analysis (ORA) on the 606 starvation-responsive genes that are dependent on *efk-1*, *zip-2*, and *cep-1* (Table S3). In agreement with the GSEA analysis, several DNA damage response and DNA repair pathways were amongst the top ORA categories, including NER and BER, two important pathways for oxidative damage repair (Figure 4F). Taken together, both functional analysis approaches suggest that *efk-1*, *zip-2*, and *cep-1* are jointly responsible for upregulating transcription of DNA repair genes during starvation.

### NER is required for starvation survival and acts in the *efk-1* pathway

NER is an evolutionarily conserved DNA repair mechanism responsible for removing bulky, DNA-distorting adducts caused by UV radiation, oxidative stress, or genotoxins (54–57). NER consists of two branches, global-genome NER (GG-NER) and transcription-coupled NER (TC-NER). In *C. elegans*, TC-NER is vital for DNA repair in somatic cells, whereas GG-NER is required in both germline and soma (55,58). To test if NER-mediated DNA repair is required for starvation survival, we assayed the starvation sensitivity of several NER mutants. We studied the *xpa-1*, *xpf-1*, *ercc-1*, and *xpg-1* mutants that are generally NER deficient, the *xpc-1* mutant that is deficient only in GG-NER, and the *csa-1* and *csb-1* mutants that are deficient only in TC-NER (59). Most NER mutants showed no visible defects in unstressed conditions, except *ercc-1*, which had a slower growth rate (Figure S5A). However, all of these mutants showed reduced survival in the L1 starvation survival assay, indicating that functional NER is required for starvation survival (Figure 5A-C). To test if NER promotes starvation survival in the *efk-1* pathway, we studied the genetic interaction between *efk-1* and *xpa-1* and *xpf-1*. *efk-1;xpa-1* and *efk-1;xpf-1* double mutants showed no genetic interaction in the L1 starvation survival assay (Figure 5D-E). Thus, functional NER is required for starvation survival in the *efk-1* pathway. We also studied genetic interactions of the NER with *zip-2* and *cep-1* by assaying *zip-2*;*xpa-1* and *zip-2*;*xpf-1* double mutants, and *cep-1;xpf-1* double mutants (we did not attempt to generate a *cep-1;xpa-1* double mutant as *cep-1* and *xpa-1* are genetically linked). In line with the above genetic interaction data, *zip-2*; *xpa-1*, *zip-2*;*xpf-1*, and *cep-1*;*xpf-1* double mutants showed no synthetic starvation survival defects (Figure 5F-H). All above double mutants had no growth defect compared to wild type (Figure S5B-F). Taken together, this supports a model whereby *efk-1*, *zip-2*, and *cep-1* jointly regulate NER to promote starvation survival.

**Figure 5.**
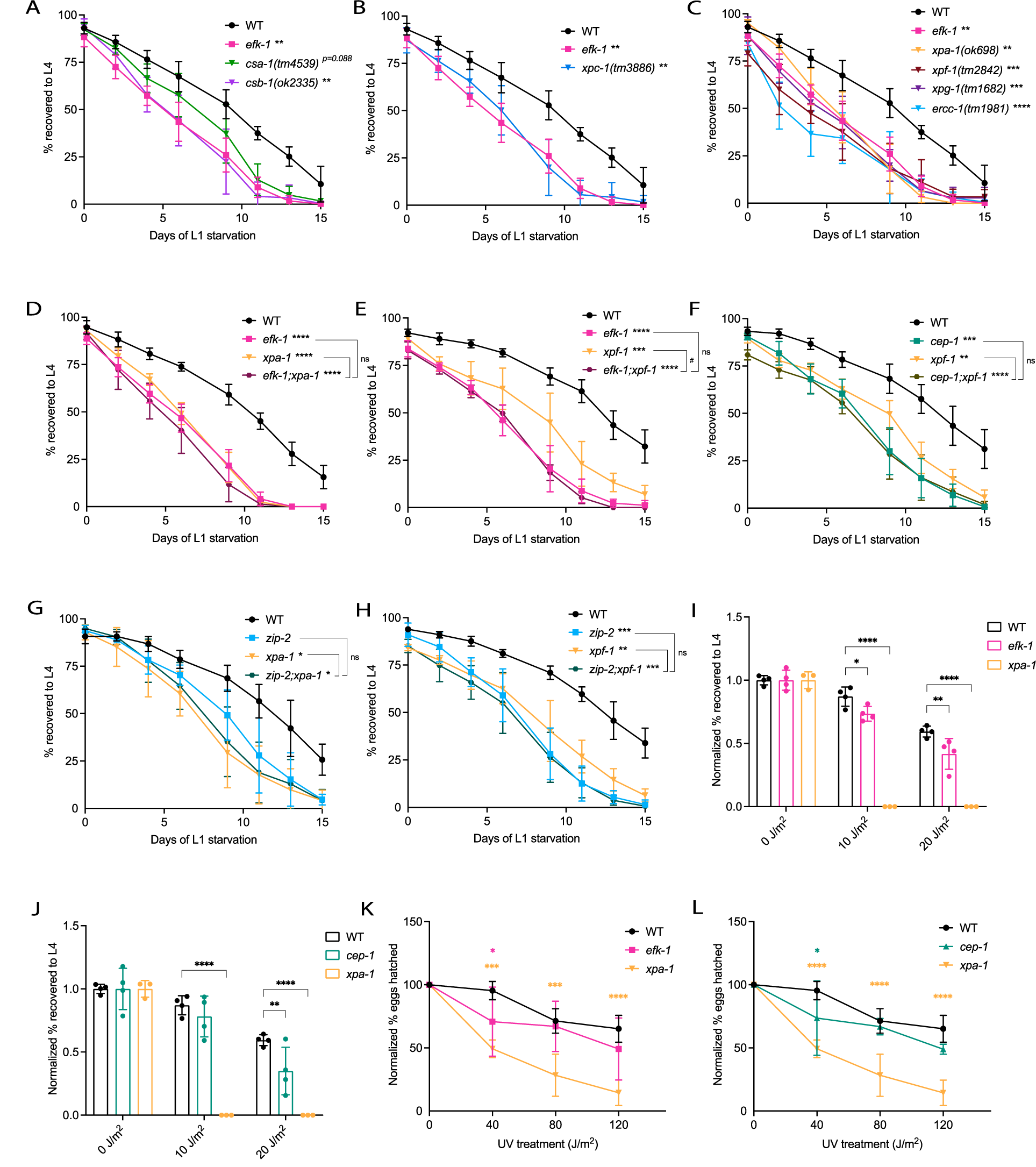
NER is required for *efk-1* mediated starvation resistance. **(A-C)** The graphs show L1 starvation survival for **(A)** TC-NER deficient *csa-1(tm4539)* and *csb-1(ok2335)* mutants, **(B)** GG-NER deficient *xpc-1(tm3886)* mutants, and **(C)** generally NER deficient *xpa-1(ok698)*, *xpf-1(tm2842)*, *xpg-1(tm1670)* and *ercc-1(tm1981)* mutants. WT and *efk-1* controls shown are of the same experiments. N=4, error bars represent SD; **p<0.01, ***p<0.001, ****p<0.0001 percent L4 vs. WT animals (AUC compared using one-way ANOVA with Dunnett’s multiple comparisons test). **(D-F)** The graphs show L1 starvation survival of **(D)** *efk-1*;*xpa-1*, **(E)** *efk-1;xpf-1*, **(F)** *cep-1;xpf-1*, **(G)** *zip-2;xpa-1*, and **(H)** *zip-2;xpf-1* double mutants alongside respective single mutants and WT control. N=4, error bars represent SD; **p<0.01, ***p<0.001, ****p<0.0001 percent L4 vs. WT animals, ^#^p<0.05 vs. *xpf-1* animals (AUC compared using one-way ANOVA with Tukey’s multiple comparisons test). **(I-J)** The graphs show UV-C larval survival of *efk-1* and *cep-1* mutants, as measured by recovery to L4 stage after UV-C irradiation (0-20 J/m^2^) at L1, normalized to no UV control. WT and *xpa-1* controls shown are of the same experiment. N=4, error bars represent SD; *p<0.05, **p<0.01, ****p<0.0001 (two-way ANOVA with Dunnett’s multiple comparisons test). **(K-L)** The graphs show UV-C induced embryo lethality of *efk-1* and *cep-1* mutants, as measured by percentages of viable embryos (24 hours post egglay) after parents were subjected to UV-C irradiation (0-120 J/m^2^) at young adult stage. WT and *xpa-1* controls shown are of the same experiment. N=4, error bars represent SD; *p<0.05, ***p<0.001, ****p<0.0001 (two-way ANOVA with Dunnett’s multiple comparisons test). WT, wild-type; ns, not significant; AUC, area under the curve. See Source data for **(A-L)**.

The soma is the main site of the starvation response during L1 arrest, as the germline only consists of two primordial germ cells that are transcriptionally quiescent (36). Somatic cells depend on TC-NER, although GG-NER is also present (58). Thus, we asked if *efk-1* acts with TC-NER, GG-NER, or both during L1 arrest, using *xpc-1* and *csb-1* mutants to assess requirements for GG- and TC-NER loss, respectively. Neither *efk-1;xpc-1* not *efk-1;csb-1* double mutants showed additive starvation survival phenotypes compared to respective single mutants (Figure S5G-I). Thus, both GG- and TC-NER are required for *efk-1*-mediated starvation resistance.

### *efk-1* and *cep-1* play a minor role in somatic and germline NER

Because they are required to induce NER genes and genetically interact with the NER pathway, we hypothesized that *efk-1*, *zip-2*, and *cep-1* may be involved in NER-mediated DNA repair in fed worms as well. To test this hypothesis, we assessed somatic and germline NER capacity by measuring UV-C sensitivity in mostly somatic L1 larvae and in the germline of young adults, respectively. To test if *efk-1*, *zip-2*, and *cep-1* are required for NER in somatic cells (58), we exposed these mutants to UV-C at the L1 stage and quantified their ability to recover to the L4 stage after irradiation. Whereas the recovery of *zip-2* mutants was similar to wild type (Figure S5J), *efk-1* and *cep-1* mutants displayed a mild reduction in their ability to recover from UV-C exposure (Figure 5I-J), indicating decreased somatic NER capacity. To test if *efk-1*, *zip-2*, and *cep-1* are required for NER in the germline (58), we exposed these mutants to UV-C as young adults, and quantified the embryonic lethality of the immediate offspring. The proportion of unhatched eggs were reduced in *efk-1* and *cep-1* mutants at 40 J/m^2^ (Figure 5K-L), whereas *zip-2* mutants showed no defect (Figure S5K), indicating that *efk-1* and *cep-1*, but not *zip-2*, are involved in germline NER. Altogether, these data show that *efk-1* and *cep-1* play a minor role in both somatic and germline NER, whereas *zip-2* is dispensable.

### BER is required for starvation survival and acts in the *efk-1* pathway

BER is a DNA repair pathway that targets oxidative base lesions, such as 8-oxoguanine (8-oxoG), 5-hydroxymethyluracil (5-hmU), and uracil (60). To test if BER-mediated DNA repair is required for starvation survival, we studied the sensitivity of BER mutants to prolonged L1 starvation. BER requires the apurinic/apyrimidinic (AP) endonucleases *apn-1* and *exo-3*, and glycosylases *ung-1* and *nth-1*, which target uracil and other oxidative lesions, respectively. We also tested the Poly ADP-Ribose Polymerase *parp-2*, which is involved in mammalian BER and conserved in *C. elegans* (60). *apn-1*, *exo-3*, *ung-1*, and *nth-1* mutants have no growth phenotypes in unstressed conditions, whereas *parp-2* mutants feature a mild developmental delay (Figure S6A). Interestingly, *apn-1*, *exo-3*, *ung-1*, *nth-1*, and *parp-2* loss-of-function mutants all showed starvation survival defects (Figure 6A). To test the interaction of *efk-1* with the BER, we constructed *efk-1;apn-1*, *efk-1;exo-3*, and *efk-1;parp-2* double mutants (Figure S6B-D). Both *efk-1;apn-1* and *efk-1;exo-3* mutants showed no additive starvation-related defects compared to respective single mutants, indicating involvement in the same pathway (Figure 6B-C). However, *efk-1;parp-2* mutants showed a substantial additive starvation survival defect (Figure 6D), suggesting that *efk-1* and *parp-2* promote starvation survival in separate pathways, with *parp-2* and *efk-1;parp-2* animals also showing slower growth in unstressed conditions (Figure S6D). Taken together, our data show that BER components are required for starvation resistance in the *efk-1* pathway, with the exception of *parp-2*.

**Figure 6.**
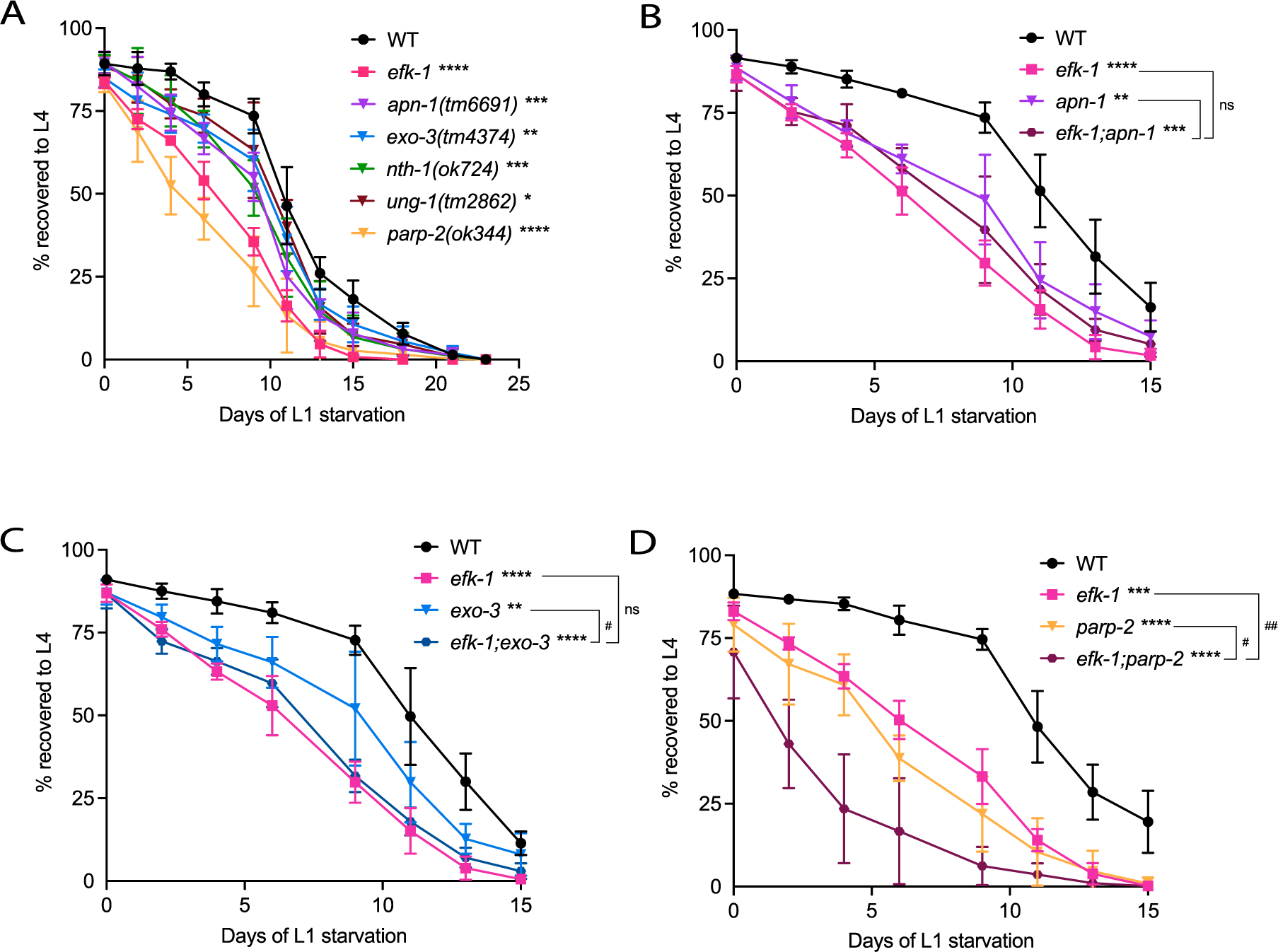
BER is required for *efk-1* mediated starvation resistance. **(A)** The graph shows L1 starvation survival for WT, *efk-1* and BER deficient *apn-1(tm6991), exo-3(tm4374), nth-1(ok724)*, *ung-1(tm2862)*, and *parp-2(ok344)* mutants. N=4, error bars represent SD; *p<0.05, **p<0.01, ***p<0.001, ****p<0.0001 percent L4 vs. WT animals (AUC compared using one-way ANOVA with Dunnett’s multiple comparisons test). **(B-D)** The graphs show L1 starvation survival of **(B)** *efk-1*;*apn-1*, **(C)** *efk-1;exo-3* and **(D)** *efk-1;parp-2* double mutants alongside respective single mutants and WT control. N=4, error bars represent SD; **p<0.01, ***p<0.001, ****p<0.0001 percent L4 vs. WT animals, ^#^p<0.05. ^##^p<0.01 between mutants (AUC compared using one-way ANOVA with Tukey’s multiple comparisons test). WT, wild-type; ns, not significant; AUC, area under the curve. See Source data for **(A-D)**.

### *efk-1* protects against elevated ROS during starvation

Oxidative stress is increased in prolonged starvation, and oxidative stress defense is required for successful starvation survival (61,62). Consistent with high oxidative burden, starving worms also accumulate more DNA lesions and express DNA damage-like signatures (63–65). To test if *efk-1* mutants have elevated ROS during starvation, we quantified ROS levels in *efk-1* mutants using two cell-permeable dyes for intracellular superoxide, dihydroethidium (DHE) and CellROX Green (61,62). Compared to wild type, *efk-1* mutants exhibited increased DHE levels from day 3 of starvation onward and increased CellROX Green levels from day 0. Importantly, both signals were rescued by the antioxidant N-acetyl cysteine (NAC; Figure 7A-B; Figure S7A-D). Thus, *efk-1* mutants accumulate abnormal ROS levels early in starvation.

**Figure 7.**
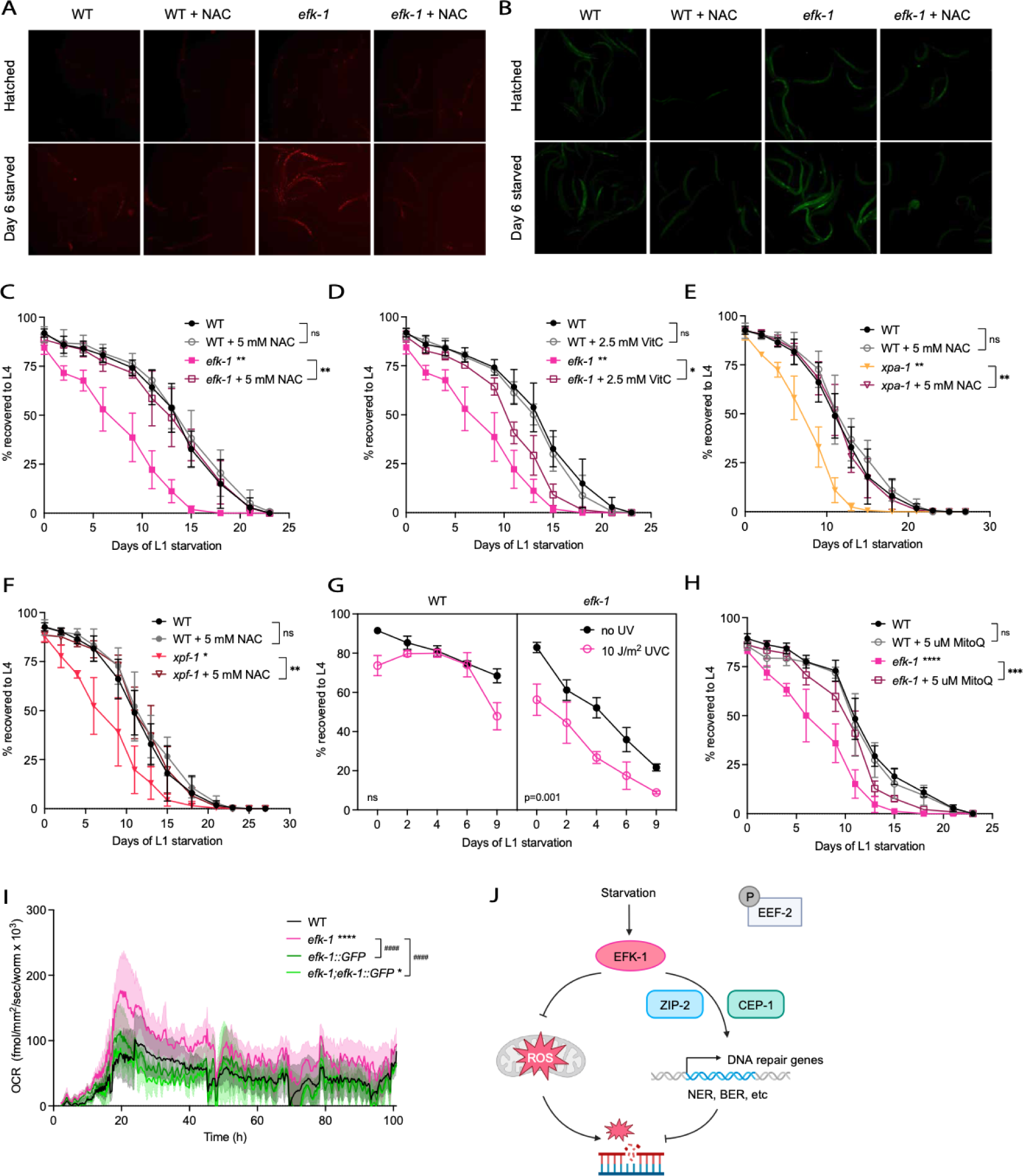
*efk-1* mutants are sensitive to starvation-induced oxidative stress, at least partially due to mitochondrial ROS. **(A-B)** The micrographs show ROS content in freshly hatched and day 6 starved WT and *efk-1* mutants with or without 5 mM NAC, as represented by superoxide dyes **(A)** dihydroethidium (DHE) and **(B)** CellROX Green. N=3 (total 100∼120 live worms per condition). **(C-D)** The graphs show L1 starvation survival of WT and *efk-1* mutants with or without supplementation of antioxidants **(C)** NAC and **(D)** VitC at 5 and 2.5 mM, respectively. WT and *efk-1* controls shown are of the same experiments. N=3, error bars represent SD; *p<0.05, **p<0.01 (AUC compared using one-way ANOVA with Tukey’s multiple comparisons test). Data are quantified in Figure S7A-D. **(E-F)** The graphs show L1 starvation survival of WT and **(E)** *xpa-1* or **(F)** *xpf-1* mutants with or without supplementation of 5 mM NAC. WT and WT+NAC controls shown are of the same experiments. N=4, error bars represent SD; *p<0.05, **p<0.01 (AUC compared using one-way ANOVA with Tukey’s multiple comparisons test). **(G)** The graph shows larval UV-C survival (y-axis) of WT and *efk-1* mutants (black, mock treatment; red, 10 J/m^2^ UVC) after pretreatment with starvation of up to 9 days (x-axis). N=3, error bars represent SD (AUC of mock treatment vs. UV-C compared using one-way ANOVA with Tukey’s multiple comparisons test). **(H)** The graph shows L1 starvation survival of WT and *efk-1* mutants with or without supplementation of mitochondrial antioxidant MitoQ at 5 uM. N=4, error bars represent SD; ***p<0.001, ****p<0.0001 (AUC compared using one-way ANOVA with Tukey’s multiple comparisons test). **(I)** The graph shows continual oxygen consumption rate (normalized OCR per worm, y-axis) of WT and *efk-1* mutants with or without the rescue construct *efk-1::GFP* in L1 starvation (hours of starvation, x-axis). N=3-4, shaded area represents standard error of the mean (SEM); *p<0.05, ****p<0.0001 vs. WT, ^####^p<0.0001 vs. *efk-1* (one-way ANOVA with Tukey’s multiple comparisons test). **(J)** Model of how *efk-1* promotes starvation survival via a noncanonical pathway. WT, wild-type; ns, not significant; ROS, reactive oxygen species; NAC, N-acetyl-L-cysteine; VitC, Vitamin C (ascorbic acid); AUC, area under the curve; MitoQ, mitoquinone; NER, nucleotide excision repair; BER, base excision repair. See Source data for **(C-I)**.

Next, we asked if oxidative stress is a cause of the starvation-induced death in the *efk-1* mutant. To address this, we treated starving L1 worms with either of two antioxidants, NAC or Vitamin C (VitC). Supplementation with 5 mM NAC completely rescued the survival defect of *efk-1* mutants (Figure 7C), whereas 2.5 mM VitC had a less substantial, but still significant effect (Figure 7D); 2.5 mM NAC and 5 mM VitC were also effective (Figure S7E-F). This shows that oxidative stress is an important cause of the starvation survival defect in *efk-1* mutants.

To test if oxidative damage is similarly the cause of the starvation-induced death in the NER mutants, we treated *xpa-1* and *xpf-1* mutants with NAC during L1 starvation. As seen with *efk-1* mutants, supplementation with 5 mM NAC completely rescued the starvation survival defects of *xpa-1* and *xpf-1* mutants (Figure 7E-F; Figure S7G-H). These data indicate that the NER promotes resistance to starvation-induced oxidative stress, likely by promoting oxidative DNA damage repair.

Next, we asked whether starvation activates the NER in an *efk-1* dependent manner. As starvation upregulates the expression of NER genes via *efk-1* (Figure 4E-F), we hypothesized that starvation would increase NER capacity and render worms more resistant to exogenous DNA damage. To test this hypothesis, we treated L1 stage wild-type and *efk-1* mutant worms with up to 9 days of starvation, then irradiated them with UV-C, and then assessed population survival. We observed that wild-type worms that have been starved for two to six days are more resistant to UV irradiation than freshly hatched (∼24h) animals or animals starved for more than six days (Figure 7G). In contrast, starved *efk-1* mutants do not show such increased resistance to UV (Figure 7G). As UV resistance is indicative of NER function (55), this increase of UV resistance post-starvation implies that starvation treatment is sufficient to activate NER in an *efk-1* dependent manner. Taken together, these data suggest that *efk-1* is required for activation of DNA repair to counteract starvation-induced oxidative DNA damage.

### *efk-1* protects against mitochondrial ROS

The mitochondria are an important site of ROS production (66), and decline of mitochondrial quality during starvation (61,63,67) correlates with oxidative stress phenotypes of starved worms. To determine if the mitochondria are a source of the increased ROS levels observed in starved *efk-1* mutants, we examined whether the starvation defect of the *efk-1* mutant was altered by the mitochondrially targeted antioxidant mitoquinone (MitoQ) (68,69), which ameliorates stress-induced mitochondrial ROS generation and protects against mitochondrial dysfunction-induced phenotypes in mammalian cells and *C. elegans* (69–71). Indeed, MitoQ supplementation partially rescued the starvation survival defect of *efk-1* mutants (Figure 7H; Figure S7I). This implies that the reduced survival of starving *efk-1* mutants is caused at least in part by mitochondrial defects, such as excess mitochondrial ROS.

To further characterize the starvation-induced mitochondrial phenotypes of *efk-1* mutants, we studied mitochondrial DNA copy number. Consistent with previous studies, we observed that mitochondrial DNA copy number decreased during L1 starvation (Figure S7J). However, mitochondrial DNA copy number in *efk-1* mutants remained similar to wild type, both in prolonged (0∼12 days) and early starvation (0∼3 days) (Figure S7J-K). Thus, altered mitochondrial content is likely not the cause of excess mitochondrial ROS in *efk-1* mutants.

Finally, we assessed respiration rates during L1 starvation by continuously monitoring oxygen consumption in live worms. Strikingly, we observed a burst of respiration in *efk-1* mutants in early starvation (16∼20 hours after hatch), which was rescued in animals expressing the *efk-1::GFP* transgene (Figure 7I; Table S4). This excess respiration cannot be attributed to increased motility, as *efk-1* mutants are no more active than wild-type worms during the specified timeframe (Figure S7L; Table S4). Thus, the excess respiration in *efk-1* mutants is more likely linked to the heightened ROS and other oxidative phenotypes, and could imply defective mitochondrial function during starvation. Together, these data show that *efk-1* likely offers protection against excess respiration and accumulation of mitochondrial ROS during L1 starvation.

## Discussion

Cells and organisms survive starvation by mounting a cellular response that involves rewiring of gene expression and metabolism. During starvation, cells reprogram gene expression from growth and development to conservation and cytoprotection. The kinase EFK-1/eEF2K is a starvation response regulator that canonically functions by allowing the cells to conserve cellular energy, specifically by attenuating translation elongation via phosphorylation of its principal substrate, EEF-2. However, emerging evidence suggests that EFK-1/eEF2K may act via additional mechanisms to promote cytoprotection. Here, we show that *C. elegans* EFK-1 acts in a noncanonical, translation elongation-independent manner to promote starvation survival. EFK-1 cooperates with TFs CEP-1/p53 and ZIP-2/bZIP to rewire transcription during early starvation, upregulating DNA repair processes such as NER and BER. In turn, these DNA repair pathways are required to counteract oxidative damage that arises from prolonged starvation. Furthermore, EFK-1 lessens oxidative burden during starvation by preventing hyperactive oxygen consumption and repressing ROS accumulation. Collectively, we propose a model whereby *efk-1* promotes starvation survival via a noncanonical pathway that involves preserving mitochondrial function, preventing accumulation of mitochondrial ROS, and promoting repair of oxidative DNA damage (Figure 7J). This is to our knowledge the first characterization of a translation-independent mechanism linked to EFK-1 mediated stress adaptation.

### *efk-1*, *cep-1*, *zip-2*, and DNA repair pathway genes define a new starvation response pathway that promotes genome integrity in starvation

*C. elegans efk-1*, *zip-2*, and *cep-1* are all required for starvation survival (5,39,41), but how each of factors promotes starvation adaptation was unknown. Our data indicate that these three regulators function in the same signaling circuit, advancing our understanding of how *C. elegans* achieves starvation survival. Inducing DNA damage response pathways is a key output of this signaling circuit, and the activity of these repair processes is itself critical for surviving starvation. Although induction of DNA repair pathways may have been expected from *cep-1*, an established guardian of genome integrity (41,72), it is a novel role for *efk-1* and *zip-2*. Additionally, we also show that *efk-1* and *cep-1* play a part in promoting DNA repair and genome stability in feeding, growing animals, whereas *zip-2* is less important in this context.

Our data provides additional links of *efk-1/eEF2K* and *cep-1/p53* in stress and DNA damage response. In cancer cells, mice, and *C. elegans*, both *cep-1*/p53 and *efk-1*/eEF2K play a similar, conserved role in mediating the cellular decision between apoptosis and DNA repair in response to genomic insults (17,49,73). In *C. elegans*, both *efk-1* and *cep-1* are required to maintain germline integrity by regulating germline apoptosis and repair (49,72,74). Consistent with these, we show that *C. elegans efk-1* and *cep-1* also act together in to promote genomic integrity in starvation. Overall, our study highlights intertwined regulatory roles of *efk-1*/*eEF2K* and *cep-1*/*p53* in evolutionarily conserved responses to genotoxic stress.

### *efk-1* protects against mitochondrial ROS

Why is it important for *C. elegans* to induce DNA repair pathways during starvation? ROS levels increase during long-term starvation in wild-type animals (61), but our data show that this phenomenon is exacerbated in *efk-1* mutants. Our data suggests that *efk-1* offers oxidative protection for the starving cell in two ways. On one hand, *efk-1* prevents excess ROS accumulation in the cell. A substantial portion of this ROS originates from the mitochondria, as shown by the ability of mitoQ to partially rescue the starvation defect in *efk-1* mutants, although non-mitochondrial ROS also play a part. On the other hand, *efk-1* protects from ROS-induced oxidative damage by activating the DNA repair machinery. In line with this model, exogenous provision of antioxidants completely rescues the starvation survival defect not only of *efk-1* mutants, but also DNA repair pathway mutants, pinpointing oxidative DNA damage as a key adverse event during starvation. However, it remains unclear how *efk-1* regulates mitochondrial physiology and function during starvation to prevent excess ROS accumulation. *efk-1* mutants display higher oxygen consumption rates, which could reflect elevated mitochondrial activity. In prolonged starvation, such increased mitochondrial burden in *efk-1* mutants might exacerbate starvation-related mitochondrial defects and cause accumulation of ROS and oxidative damage in the cell.

### *efk-1* regulates a translation-independent pathway

Mammalian eEF2K protects against starvation via inducible eEF2 phosphorylation and translation attenuation. However, we found contrary evidence suggesting that this is not a mechanism of *efk-1*-mediated starvation resistance in *C. elegans*. Although EFK-1 is also the sole kinase for EEF-2 phosphorylation in *C. elegans* (16,17), substantial EEF-2 phosphorylation is present in feeding worms and appears unperturbed by standard starvation treatments, regardless of duration of starvation or animal developmental stage (Figure 2A-C). These observations suggest that EFK-1 is constitutively active in *C. elegans* fed a standard *E. coli* diet. Interestingly, EFK-1 overexpression, measured to approximate a 10-fold increase in mRNA, did not increase p-EEF-2 levels past the observed baseline. It is possible that EFK-1 kinase is constitutively active in *C. elegans*, such that EEF-2 phosphorylation is saturated.

Our data show that *efk-1* exerts influence on gene expression via rewiring transcription, although it is unclear how *efk-1* signals to these transcription factors. We did not observe an *efk-1*-dependent increase in *zip-2* and *cep-1* transcript in starved worms (Table S1), suggesting that their activity may be regulated on a translational or post-translational level. As eEF2K phosphorylates substrates other than EEF-2 (15), it is possible that EFK-1 could signal to ZIP-2 and CEP-1 directly via phosphorylation. Alternatively, EFK-1 may regulate the expression of downstream factors via selective translation without affecting the global translation rate, which has been observed in mammalian cells and *C. elegans* under physiological stress conditions (1,75). EFK-1 may employ either or both of these modes of regulation to activate the downstream stress response pathway. These possible new mechanisms of *efk-1*/eEF2K-mediated stress signaling may be explored in future studies.

Taken together, our evidence points to a noncanonical, translation-independent mechanism of EFK-1 in *C. elegans* in starvation adaptation. To date, most studies on eEF2K have been conducted in transformed mammalian cell lines, which are fundamentally different from whole organisms such as *C. elegans*. It is unknown why *C. elegans efk-1* employs a distinct mechanism, whether due to evolutionary divergence, differences between transformed and non-transformed cells, or tissue-specific effects. Our study has uncovered hitherto unknown regulatory roles of EFK-1 that may prove valuable to understanding how eEF2K regulates stress responses in different biological and evolutionary contexts.

## Methods

### *C. elegans* strains and maintenance

We cultured *C. elegans* strains using standard techniques on nematode growth media-lite (NGM-l) plates (76). Each mutant was crossed into our lab N2 strain at least six times to remove background mutations. To prevent accumulation of germline mutations, DNA repair mutants were frozen immediately after the final backcross, and an aliquot was thawed and maintained for no more than four generations prior to each experiment. *E. coli* OP50 was the standard food source; HT115 was used for RNAi experiments, and streptomycin resistant OP50-1 was used for the starvation-primed UV recovery assay. All experiments were carried out at 20°C unless otherwise specified. Worm strains used in this study are listed in the Key Resource Table.

For synchronized worm growths, we isolated embryos by standard sodium hypochlorite treatment, and residual hypochlorite was removed by washing 3 times in M9 (77). Isolated embryos were kept overnight in sterile M9 or on unseeded NGM-l plates until all hatched and arrested at the L1 stage via short-term fasting (16–24 hr). Synchronized L1 stage larvae were then transferred to seeded plates and grown to the desired stage.

Standard genetic crossing techniques were used to construct double mutants; genotyping primers are indicated in the Key Resource Table. We attempted to construct the *efk-1(ok3609);cebp-2(tm5421)* double mutant, but failed to find homozygous double mutant animals from the clonal progeny of 100 individual candidate F2 worms, suggesting that loss of both genes causes synthetic lethality.

Feeding RNAi was performed on NGM-l plates supplemented with 25 μg/ml carbenicillin (BioBasic CDJ469), 1 mM IPTG (Santa Cruz CAS 367-93-1), and 12.5 μg/ml tetracycline (BioBasic TB0504; NGM-l-RNAi plates), and seeded with appropriate HT115 RNAi bacteria. The RNAi clones were from the Ahringer library (Source BioScience) and were sequenced prior to use.

### Generation of transgenic strains

To generate the transgenic *efk-1p::EFK-1::GFP* reporter strain, we used Phusion Hot Start II DNA Polymerase (Thermo Scientific F549L) to amplify 2000 bp upstream of *efk-1* start codon, and the *efk-1* genomic sequence including introns but without the final stop codon. *efk-1* promoter and genomic sequence were cloned into Zero Blunt™ TOPO™ vector (Invitrogen 450245), transformed into TOPO One Shot™ MAX Efficiency DH5α-T1^R^ *E. coli* (Invitrogen 12297016), followed by restriction digestion and insertion into pSM vector (78) between restriction sites HindIII / AscI, and AscI / KpnI, respectively. 50 ng/ul of each plasmid and 25 ng/μl of co-injection marker *Podr-1::RFP* were microinjected into the gonad of day-one adult worms, and F2 were screened for stable transmission. We found a random integrant at 100% transmission and validated that both GFP and marker segregated in Mendelian proportions over three generations.

### *C. elegans* starvation studies

Worms were maintained for at least two generations in a fed state prior to all starvation experiments. For L4 studies, we synchronized worms by hatching in M9 overnight (16∼24 hours), followed by growth on seeded 100mm NGM-l plates to mid-L4 at a density of 2000-2200 worms/plate. The fed portion was collected at mid L4 stage as judged by vulval morphology developmental staging. The starved portion was first washed 5 times in sterile M9, and then placed either on unseeded plates or suspended in sterile M9 for the number of hours specified.

For L1 studies, worms were synchronized and grown on seeded plates to gravid adulthood at a density of 1000-1200 worms/plate. Worms were washed off the plate and residual embryos were collected with a cell scraper (Falcon #353085). For Western blot and RT-qPCR studies, bulk embryos were isolated by vigorous mixing in hypochlorite solution (40 ml dH2O, 7 ml sodium hypochlorite Sigma #425044, 3 ml 1M KOH) for 6-7 minutes, hatched overnight without food, and subjected to starvation treatments as specified. For imaging and mitochondrial DNA content studies, worms were bleached for 2 minutes, and excess carcass was removed by filtering through a 35-micron nylon mesh (Elko Filtering #03-35/16).

L1 starvation survival assays were adapted from (79). Synchronized embryos were extracted by hypochlorite treatment for 2 minutes and resuspended in S-basal medium without cholesterol at a concentration of ∼1 worm/μl and supplemented with an antibiotic–antimycotic mix (Gibco #15240062). Worms were starved at 20°C while being continuously rotated on a tube rotator (ThermoFisher) at 20 rpm. To assess viability, ∼150 animals were transferred to seeded NGM-l plates every 2-3 days and assessed for growth to the L4 stage after 48h at 20°C, with day 0 being counted as 16-24h after hypochlorite treatment. Statistical significance of three to four independent biological repeats was calculated using area under the curve and one-way ANOVA with Tukey’s or Dunnett’s multiple comparisons test.

For starvation studies with chemical treatments, CHX (Sigma #1810), puromycin (Sigma #P8833), NAC (Sigma #A7250), VitC (Sigma #A92902), and mitoQ (MedChemExpress #HY-100116A) stock solutions were prepared freshly in S-basal at concentrations of 10 mM, 10 mM, 100 mM, 100 mM, and 5 mM, respectively. Each chemical was filtered through a 0.2 um filter and added to the culture at specified concentrations. To compensate for degradation of NAC in aqueous solution during L1 starvation survival experiments (80), we added fresh NAC at 40% of the original amount on day 6. VitC containing solutions were kept in the dark in aluminum foil to minimize photodegradation (81).

### A549 starvation studies

A549 cells were obtained from Dr. Poul Sorenson. For Western blot experiments, 2 x 10^5^ cells were seeded in 6-well plates and grown overnight in Dulbecco’s modified Eagle’s medium (DMEM) (Gibco #11995073) supplemented with 10% fetal bovine serum (FBS) (Gibco #12483020) at 37°C under 5% CO_2_ in a humidified incubator. The media was aspirated, and the fed condition was incubated in 10% FBS DMEM, while the starved condition was incubated in Hanks’ Balanced Salt Solution (HBSS) (Gibco #14175095) for 3, 6, and 24 hours. Total cellular protein was extracted using RIPA lysis buffer supplemented with protease inhibitor and phosphatase inhibitors.

### Growth rate assays

We measured the growth rate for every strain used in the L1 starvation survival assay. Staged L4 worms were picked onto a seeded plate and grown to gravid adults overnight at RT. 10 gravid adults were picked onto a fresh plate and allowed to lay eggs for 2h at RT. Parent animals were removed and synchronized F1 were incubated in 20°C for 56h. The portion of F1 that were able to reach L4 stage was recorded. Statistical analysis was performed using one-way ANOVA with Dunnett’s multiple comparisons test.

### UV sensitivity assays

UV sensitivity assays were adapted from (58,82). For the somatic NER assay, L1 worms were synchronized overnight on unseeded NGM-l plates, then irradiated uncovered in a Stratalinker 2400 UV Crosslinker (Stratagene) with wavelength 254 nm light at specified dosages. Worms were placed at 20°C for 48h and scored for recovery to L4 stage. Recovery rates were normalized to no UV control of the same genotype. Statistical analysis was performed using two-way ANOVA with Dunnett’s multiple comparisons test.

For the germline NER assay, embryos were extracted by hypochlorite treatment and placed on seeded NGM-l plates at 20°C for 72 hours until young adulthood. Worms were collected, washed three times in M9, and placed on unseeded NGM-l plates. Worms were allowed to disperse for an hour and then irradiated uncovered in the Stratalinker 2400 UV Crosslinker at specified dosages. Immediately following UV exposure, 15-20 staged young adults were picked onto a seeded NGM-l plate and allowed to lay eggs for 4 hours. The proportion of eggs that hatched after 24 hours was recorded. Embryonic lethality was normalized to no UV control of the same genotype. For each dosage, statistical significance between mutants was computed using two-way ANOVA with Dunnett’s multiple comparisons test.

For testing NER activation after exposure to starvation, worms were starved in S-basal for the specified number of days as described above. At each timepoint, a portion of worms were placed on an unseeded 60 mm plate and allowed to disperse for an hour. Uncovered plates were exposed to UV as above. After irradiation, worms were quantified, placed on a seeded OP50-1 plate for recovery at 20°C for 48 hours, and scored for recovery to L4 stage. Statistical significance of three independent biological repeats was calculated using area under the curve and one-way ANOVA with Tukey’s multiple comparisons test.

### Respiration and motility measurement

Measurement of oxygen consumption rates (OCR) in live *C. elegans* was conducted with the Resipher system (Lucid Scientific) as described (83). Embryos from wild-type, *efk-1(ok3609)*, *efk-1::GFP*, and *efk-1(ok3609);efk-1::GFP* worms were isolated by hypochlorite treatment and placed on bacteria-free NGM plates for L1 synchronization. A 96-well microplate was seeded with synchronized L1 worms at a density of 100 worms in 100 µL of M9 buffer per well and fitted with the Resipher sensor lid. The Resipher device continuously recorded OCR data for 101 hours with timepoints taken every 15 minutes. Afterwards, OCR rates in each well were normalized to the number of worms in that well as determined by visual inspection. OCR data normalized per worm were averaged across technical replicates (6-12 wells) for each time point in each of 3-4 biological replicates. Time-course data for the 3-4 biological replicates was analyzed by a global area under the curve (AUC) analysis in GraphPad Prism 10. The computed AUC values and associated standard errors were compared across the four groups using one-way ANOVA with Tukey’s multiple comparison test.

*C. elegans* motility was recorded continuously for 40 hours with the wMicrotracker (Phylum Tech). Wild-type, *efk-1(ok3609)*, *efk-1::GFP*, and *efk-1(ok3609);efk-1::GFP* embryos were isolated and synchronized as for the Resipher analysis. A 96-well microplate was seeded with synchronized L1 worms at a density of 100 worms in 100 µL of M9 buffer in each well. Movement data was recorded every hour. Afterwards, motility rates were normalized to the number of worms present in each well as determined by visual inspection. Similar to the Resipher analysis, motility data was normalized per worm were averaged across technical replicates (6-12 wells) for each time point in each of 3 biological replicates. Statistical analysis was performed as for the Resipher data.

### SDS-PAGE and Western blotting

For *C. elegans* studies, synchronized mid-L4 worms or L1 worms were collected, rapidly washed once with M9 to remove excess bacteria, and flash frozen in ethanol and dry ice. Worm pellets were sonicated in 1 ml ice-cold Protein Lysis Buffer (150 mM KCl, 1 mM EDTA, 0.25% SDS, 1.0% NP-40, 50 mM Tris/HCl pH=7.4) supplemented with 1 mM PMSF, 1x Roche cOmplete™ Protease Inhibitor Cocktail (#4693116001, Roche), 50 mM NaF and 1 mM sodium orthovanadate. For A549 cell studies, cells were washed by cold phosphate-buffered saline (PBS) twice and lysed with 400 ul RIPA buffer (150 mM NaCl, 50 mM Tris/HCl pH=7.4, 1.0% NP-40, 0.5% DOC, 0.1% SDS) supplemented with 1.15 mM PMSF, 1 x Roche cOmplete^TM^ Protease Inhibitor Cocktail, 50 mM NaF, and 1mM sodium orthovanadate. Protein concentrations were determined using the RC DC Protein Assay kit (#500-0121, Bio-Rad) against a BSA standard curve, and 5 ug of protein was loaded in each well. SDS-PAGE analysis and immunoblotting were performed as described (40). For *C. elegans* studies, primary antibodies were diluted in 2.5% skimmed milk as follows: 1:1000 anti-rabbit phospho-eEF2 (Thr56) antibody (#2331, Cell Signaling Technologies), 1:1000 anti-rabbit eEF2 antibody (#2332, Cell Signaling Technologies), 1:500 anti-rabbit phospho-eIF2α (Ser51) antibody (#9721, Cell Signaling Technologies), 1:1000 anti-mouse GFP antibody (#11814460001, Roche), 1:1000 anti-mouse α-tubulin antibody (#T9026, Sigma), or 1:1000 anti-mouse puromycin antibody (#MABE343, Millipore). For A549 cell line studies, primary antibodies were diluted in 5% skimmed milk as above, except for tubulin which was used at 1:5000 concentration. Anti-rabbit (#7074, NEB) and anti-mouse (#7076, Cell Signaling Technologies) HRP conjugated secondary antibodies were diluted at 1:5000 in 5% skimmed milk. Detection was done using ECL (#32109, Pierce) in a Vilber Fusion FX chemiluminescence imager. Quantification of relative signal intensity was done using ImageJ. Mean value of each band was measured with a selection of the same area, inverted, and corrected for background. Relative signal intensity was obtained by dividing the background-corrected intensity of the protein bands with that of the tubulin loading control or EEF-2, as appropriate. Statistical significance was calculated using one-way ANOVA or two-way ANOVA with the statistical test specified in Source Data.

### Puromycin incorporation assay (SUnSET)

SUnSET was adapted from previous studies (26,32). For *C. elegans* studies, synchronized L4 or L1 worms were starved for the time specified. CHX controls were treated with 2 mg/ml CHX (Sigma #1810) added 6 hours prior to harvest. All samples were incubated with 0.5 mg/ml puromycin for 3 hours before harvest. Worm pellets were washed once in M9 and flash frozen. For A549 cell line studies, 2 x 10^5^ cells were fed or starved 6 hours as above. CHX controls were treated with 2 ug/ml CHX 1 hour before harvest. All samples were supplemented with 1 ug/ml puromycin 10 mins before harvest. Statistical significance was calculated using two-way ANOVA with uncorrected Fisher’s LSD.

### DIC and fluorescence microscopy

For *efk-1::GFP* images, synchronized mid-L4 *efk-1::GFP* worms were immobilized in 12.5 mM levamisole (Sigma L9756) on 2% (w/v) agarose pads. Images were captured at 40x magnification on a Hamamatsu ORCA-Flash4.0 LT+ Digital CMOS camera attached to a Leica SP8X confocal microscope.

To analyze fluorescence in the *irg-1p::GFP* reporter line, synchronized L1s were placed onto NGM-l plates seeded with OP50 or RNAi plates seeded with the appropriate HT115 RNAi culture. *irg-1p::GFP* worms were kept at 20°C for 48h until they reached mid L4-stage. *fmo-2p::GFP* worms were kept at 22°C for 48h until they reached young adulthood. Worms were immobilized as above, and images were captured at 10x magnification on a CoolSnap HQ camera (Photometrics) attached to a Zeiss Axioplan 2 compound microscope. Images were captured at 300 ms exposure time for GFP.

### ROS staining

Synchronized L1 worms were starved in S-basal as above. At each timepoint of starvation, an aliquot was treated with either DHE or CellROX Green, as adapted from (61,62). For DHE, worms were washed 3 times in M9, incubated with 10 μM DHE (Sigma #D7008) for 30 min on a tube rotator, washed 3 times with PBS, and mounted onto 2% agarose pads in 12.5 mM levamisole for imaging. For CellROX Green, worms were washed 3 times in M9, fixed in 2% formaldehyde (Sigma #F8775) for 30 min, washed 3 times with M9 with 0.01% Tween, incubated with 5 μM CellROX Green for one hour, washed 3 times with PBS and mounted onto 2% agarose pads for imaging. L1 worms were imaged using the 20X objective. Images were captured at 300 ms exposure time for CellROX Green and 100 ms for DHE. For each experiment, 3 independent biological repeats were performed with 90∼110 live worms quantified per timepoint. Images were analyzed using ImageJ software (https://imagej.nih.gov/ij/download.html). Normalized whole worm fluorescence was quantified by measuring the whole-body fluorescence of worms minus the average of 3 spots of background taken on the same image. For each timepoint, statistical significance between conditions was calculated using one-way ANOVA with Tukey’s multiple comparisons test.

### RNA sequencing and GO term analysis

Synchronized L1 wild-type, *efk-1(ok3609)*, *zip-2(tm4248)*, and *cep-1(gk138)* worms were allowed to grow on OP50 plates to mid-L4 stage. Fed samples was collected immediately by flash-freezing. Starved samples were washed 5 times in sterile M9, starved for 8 hours on unseeded plates, and then collected. RNA was isolated from whole worms as described (84). RNA integrity and quality were ascertained on a BioAnalyzer. Construction of strand-specific mRNA sequencing libraries and sequencing (75bp PET) on an Illumina HiSeq 2500 machine was done at the Sequencing Services facility of the Genome Sciences Centre, BC Cancer Agency, Vancouver BC, Canada (https://www.bcgsc.ca/services/sequencing-services). We sequenced >20 million reads per sample. The raw FASTQ reads obtained from the facility were trimmed using Trimmomatic version 0.36 (85) with parameters LEADING:3 TRAILING:3 SLIDINGWINDOW:4:15 MINLEN:36. Next, the trimmed reads were aligned to the NCBI reference genome WBcel235 WS277 (https://www.ncbi.nlm.nih.gov/assembly/GCF_000002985.6/) using Salmon version 0.9.1 (86) with parameters -l A -p 8 --gcBias. Then, transcript-level read counts were imported into R and summed into gene-level read counts using tximport (87). Genes not expressed at a level greater than 1 count per million (CPM) reads in at least three of the samples were excluded from further analysis. The gene-level read counts were normalized using the trimmed mean of M-values (TMM) in edgeR (88) to adjust samples for differences in library size. Differential expression analysis was performed using the quasi-likelihood F-test with the generalized linear model (GLM) approach in edgeR (88). RNA-seq data have been deposited at NCBI Gene Expression Omnibus (https://www.ncbi.nlm.nih.gov/geo/) under the record GSE259223. Functional enrichment analysis and visualization were performed using easyGSEA and easyVizR in the eVITTA toolbox (https://tau.cmmt.ubc.ca/eVITTA/; input September 19, 2023) (89).

For the reanalysis of published starvation RNA-seq data (20), raw reads were downloaded from Sequence Read Archive (SRA) accession PRJNA386884, extracted using fastq-dump, and processed as above. Differential expression analysis was performed as described above. Venn diagrams of gene set overlaps and associated p-value was generated as described (90).

### RNA isolation and qRT-PCR analysis

RNA isolation was performed as previously described (84). 2 μg total RNA was used to generate cDNA with Superscript II reverse transcriptase (Invitrogen 18064-014), random primers (Invitrogen 48190-011), dNTPs (Fermentas R0186), and RNAseOUT (Invitrogen 10777-019).

Quantitative real-time PCR was performed in 10 μl reactions using PowerTrack™ SYBR Green Master Mix (Applied Biosystems A46109), 1:10 diluted cDNA, and 5 μM primer, and analyzed with an Applied Biosystems QuantStudio 5 Real-Time PCR System. We analyzed the data with the ΔΔCt method. For each sample, we calculated normalization factors by averaging the (sample expression)/(average wild type expression) ratios of three normalization genes, *act-1*, *tba-1*, and *ubc-2*. Statistical significance was calculated using unpaired t-test. All data originate from three or more independent biological repeats, and each PCR reaction was conducted in technical triplicate. RT-qPCR Primers were tested on serial cDNA dilutions and analyzed for PCR efficiency prior to use. Sequences of qRT-PCR primers are listed in the Key Resource Table.

### Mitochondrial DNA content assay

Mitochondrial DNA content was measured using a qPCR-based assay as described (91). Synchronized L1 worms were starved as above. For each timepoint, 300 worms were lysed in 30 ul lysis buffer as described, and 1 ul of 1:10 diluted lysate was used per 10 ul qPCR reaction performed in quadruplicates. Statistical analysis was performed using two-way ANOVA with Tukey’s multiple comparisons test. Primers that amplify mtDNA and a noncoding nuclear region were taken from (91) and listed in Key Resource Table.

## Data availability

The authors declare that the data underlying the findings of this study are available within the paper and its Supplementary Information files. Source data underlying the Figures and Supplementary Figures are provided in a Source Data file. RNA-seq transcriptomics datasets are available at Gene Expression Omnibus (GEO accession GSE259223).

## Acknowledgements

We thank Dr. E. R. Troemel (UCSD) and Dr. Alberto Delaidelli (BC Cancer Agency) for providing invaluable suggestions and advice, and Taubert lab members for comments on the manuscript. We thank Rachel Cheng for generating *C. elegans* strains, Dr. Kota Mizumoto (UBC) for providing cloning vector pSM, Dr. E. R. Troemel for providing backcrossed *zip-2* and *cebp-2* strains, and Dr. N. J. O’Neil for providing backcrossed *xpa-1*, *ercc-1*, and *xpf-1* mutant strains. Some strains were provided by the CGC, which is funded by NIH Office of Research Infrastructure Programs (P40 OD010440). Grant support was from The Canadian Institutes of Health Research (CIHR; PJT-153199 and PJT-186144 to ST; FDN-143280 to PHS), the Natural Sciences and Engineering Research Council of Canada (NSERC; RGPIN-2018-05133 to ST), the British Columbia Cancer Foundation through generous donations from Team Finn and other riders in the Ride to Conquer Cancer, and a research grant from the National Institutes of Health (R00 ES029552 to JHH). JY was supported by BCCHR and UBC scholarships, CS by a BCCHR scholarship, PB by an NSERC undergraduate student research award (USRA), DS by a UBC Faculty of Medicine Summer Student Research Program (FoM SSRP) award, and ST by a Canada Research Chair and a BCCHR Investigator Grant Award Program (IGAP) award.

## Author contributions

**Design of research:** JY, FB, CS, JNM, PHS, JHH, ST. **Performed experiments:** JY, FB, TTM, CS, PB, GD, DS, JHH, ST. **Analysed data:** JY, FB, XC, TTM, JHH, ST. **Wrote the manuscript:** JY, ST. **Edited the manuscript:** All authors.

---

| **<u>Reagent type (species) or resource</u>** | **<u>Designation</u>** | **<u>Source or reference</u>** |
| --- | --- | --- |
| Strain, strain background (*Escherichia coli*) | OP50 | Caenorhabditis Genetics Center (CGC) |
| Strain, strain background (*Escherichia coli*) | OP50-1 | Caenorhabditis Genetics Center (CGC) |
| Strain, strain background (*Escherichia coli*) | HT115 | Caenorhabditis Genetics Center (CGC) |
| Strain, competent cells (*Escherichia coli*) | DH5α Competent Cells | Thermo Scientific™ |
| Strain, competent cells (*Escherichia coli*) | TOP10 Competent Cells | Thermo Scientific™ |
| genetic reagent (*C. elegans*) | N2 | Caenorhabditis Genetics Center (CGC) |
| genetic reagent (*C. elegans*) | efk-1(ok3609) III | Caenorhabditis Genetics Center (CGC) |
| genetic reagent (*C. elegans*) | steIs57 [efk-1p::EFK-1::GFP + Podr-1::RFP] | This paper |
| genetic reagent (*C. elegans*) | efk-1(ok3609) III; steIs57 [efk-1p::EFK-1::GFP + Podr-1::RFP] | This paper |
| genetic reagent (*C. elegans*) | agIs17 [myo-2p::mCherry + irg-1p::GFP] IV. | Caenorhabditis Genetics Center (CGC) |
| genetic reagent (*C. elegans*) | zip-2(tm4248) III | Dr. E. Troemel (UCSD) |
| genetic reagent (*C. elegans*) | cebp-2(tm5421) I | Dr. E. Troemel (UCSD) |
| genetic reagent (*C. elegans*) | cep-1(gk138) I | Caenorhabditis Genetics Center (CGC) |
| genetic reagent (*C. elegans*) | xpa-1(ok698) I | Dr. N. J. O'Neil (UBC) |
| genetic reagent (*C. elegans*) | xpf-1(tm2842) II | Dr. N. J. O'Neil (UBC) |
| genetic reagent (*C. elegans*) | csa-1(tm4539) II | NBRP |
| genetic reagent (*C. elegans*) | csb-1(ok2335) X | Caenorhabditis Genetics Center (CGC) |
| genetic reagent (*C. elegans*) | xpc-1(tm3886) IV | NBRP |
| genetic reagent (*C. elegans*) | ercc-1(tm1981) I | Dr. N. J. O'Neil (UBC) |
| genetic reagent (*C. elegans*) | xpg-1(tm1682) I | NBRP |
| genetic reagent (*C. elegans*) | efk-1(ok3609) III; xpa-1(ok698) I | This paper |
| genetic reagent (*C. elegans*) | efk-1(ok3609) III; xpf-1(tm2842) II | This paper |
| genetic reagent (*C. elegans*) | zip-2(tm4248) III; xpa-1(ok698) I | This paper |
| genetic reagent (*C. elegans*) | zip-2(tm4248) III; xpf-1(tm2842) II | This paper |
| genetic reagent (*C. elegans*) | cep-1(gk138) I; xpf-1(tm2842) II | This paper |
| genetic reagent (*C. elegans*) | efk-1(ok3609) III; xpc-1(tm3886) IV | This paper |
| genetic reagent (*C. elegans*) | efk-1(ok3609) III; csb-1(ok2335) X | This paper |
| genetic reagent (*C. elegans*) | apn-1(tm6691) II | NBRP |
| genetic reagent (*C. elegans*) | exo-3(tm4374) I | NBRP |
| genetic reagent (*C. elegans*) | nth-1(ok724) III | Caenorhabditis Genetics Center (CGC) |
| genetic reagent (*C. elegans*) | ung-1(tm2862) III | Caenorhabditis Genetics Center (CGC) |
| genetic reagent (*C. elegans*) | parp-2(ok344) II | Caenorhabditis Genetics Center (CGC) |
| genetic reagent (*C. elegans*) | efk-1(ok3609) III; apn-1(tm6691) II | This paper |
| genetic reagent (*C. elegans*) | efk-1(ok3609) III; exo-3(tm4374) I | This paper |
| genetic reagent (*C. elegans*) | efk-1(ok3609) III; parp-2(ok344) II | This paper |
| Sequence-based reagent | Fwd_HindIII_efk-1P | This paper |
| Sequence-based reagent | efk-1p_pSM_SgsI_rev | This paper |
| Sequence-based reagent | efk-1_pSM_SgsI_fwd | This paper |
| Sequence-based reagent | efk-1_pSM_KpnI_rev | This paper |
| Sequence-based reagent | efk-1_outer_fwd | This paper |
| Sequence-based reagent | efk-1_outer_rev | This paper |
| Sequence-based reagent | TOPO_M13_fwd | Zero Blunt® TOPO® PCR Cloning Kit |
| Sequence-based reagent | TOPO_M13_rev | Zero Blunt® TOPO® PCR Cloning Kit |
| Sequence-based reagent | efk-1p_TOPO_seq2f | This paper |
| Sequence-based reagent | efk-1_TOPO_seq2f | This paper |
| Sequence-based reagent | efk-1_TOPO_seq3r | This paper |
| Sequence-based reagent | efk-1_TOPO_seq4f | This paper |
| Sequence-based reagent | pSM_seq_fwd | This paper |
| Sequence-based reagent | pSM_seq_rev | This paper |
| Sequence-based reagent | mtDNAqPCR_mito_fwd | https://www.ncbi.nlm.nih.gov/pmc/articles/PMC7817664/ |
| Sequence-based reagent | mtDNAqPCR_mito_rev | https://www.ncbi.nlm.nih.gov/pmc/articles/PMC7817664/ |
| Sequence-based reagent | mtDNAqPCR_ctrl_ges-1_fwd | https://www.ncbi.nlm.nih.gov/pmc/articles/PMC7817664/ |
| Sequence-based reagent | mtDNAqPCR_ctrl_ges-1_rev | https://www.ncbi.nlm.nih.gov/pmc/articles/PMC7817664/ |
| Sequence-based reagent | efk-1_ext_fwd | This paper |
| Sequence-based reagent | efk-1_ext_rev | This paper |
| Sequence-based reagent | efk-1_int_fwd | This paper |
| Sequence-based reagent | efk-1_int_rev | This paper |
| Sequence-based reagent | zip-2_ext_fwd | This paper |
| Sequence-based reagent | zip-2_ext_rev | This paper |
| Sequence-based reagent | zip-2_int_fwd | This paper |
| Sequence-based reagent | zip-2_int_rev | This paper |
| Sequence-based reagent | cep-1_ext_fwd | This paper |
| Sequence-based reagent | cep-1_ext_rev | This paper |
| Sequence-based reagent | cep-1_int_fwd | This paper |
| Sequence-based reagent | cep-1_int_rev | This paper |
| Sequence-based reagent | xpa-1_ext_fwd | This paper |
| Sequence-based reagent | xpa-1_ext_rev | This paper |
| Sequence-based reagent | xpa-1_int_fwd | This paper |
| Sequence-based reagent | xpa-1_int_rev | This paper |
| Sequence-based reagent | xpf-1_ext_fwd | This paper |
| Sequence-based reagent | xpf-1_ext_rev | This paper |
| Sequence-based reagent | xpf-1_int_fwd | This paper |
| Sequence-based reagent | xpf-1_int_rev | This paper |
| Sequence-based reagent | csa-1_ext_fwd | This paper |
| Sequence-based reagent | csa-1_ext_rev | This paper |
| Sequence-based reagent | csa-1_int_fwd | This paper |
| Sequence-based reagent | csa-1_int_rev | This paper |
| Sequence-based reagent | csb-1_ext_fwd | This paper |
| Sequence-based reagent | csb-1_ext_rev | This paper |
| Sequence-based reagent | csb-1_int_fwd | This paper |
| Sequence-based reagent | csb-1_int_rev | This paper |
| Sequence-based reagent | xpc-1_ext_fwd | This paper |
| Sequence-based reagent | xpc-1_ext_rev | This paper |
| Sequence-based reagent | xpc-1_int_fwd | This paper |
| Sequence-based reagent | xpc-1_int_rev | This paper |
| Sequence-based reagent | ercc-1_ext_fwd | This paper |
| Sequence-based reagent | ercc-1_ext_rev | This paper |
| Sequence-based reagent | ercc-1_int_fwd | This paper |
| Sequence-based reagent | ercc-1_int_rev | This paper |
| Sequence-based reagent | xpg-1_ext_fwd | This paper |
| Sequence-based reagent | xpg-1_ext_rev | This paper |
| Sequence-based reagent | xpg-1_int_fwd | This paper |
| Sequence-based reagent | xpg-1_int_rev | This paper |
| Sequence-based reagent | apn-1_ext_fwd | This paper |
| Sequence-based reagent | apn-1_ext_rev | This paper |
| Sequence-based reagent | apn-1_int_fwd | This paper |
| Sequence-based reagent | apn-1_int_rev | This paper |
| Sequence-based reagent | exo-3_ext_fwd | This paper |
| Sequence-based reagent | exo-3_ext_rev | This paper |
| Sequence-based reagent | exo-3_int_fwd | This paper |
| Sequence-based reagent | exo-3_int_rev | This paper |
| Sequence-based reagent | nth-1_ext_fwd | This paper |
| Sequence-based reagent | nth-1_ext_rev | This paper |
| Sequence-based reagent | nth-1_int_fwd | This paper |
| Sequence-based reagent | nth-1_int_rev | This paper |
| Sequence-based reagent | ung-1_ext_fwd | This paper |
| Sequence-based reagent | ung-1_ext_rev | This paper |
| Sequence-based reagent | ung-1_int_fwd | This paper |
| Sequence-based reagent | ung-1_int_rev | This paper |
| Sequence-based reagent | parp-2_ext_fwd | This paper |
| Sequence-based reagent | parp-2_ext_rev | This paper |
| Sequence-based reagent | parp-2_int_fwd | This paper |
| Sequence-based reagent | parp-2_int_rev | This paper |
| Sequence-based reagent | efk-1_F | This paper |
| Sequence-based reagent | efk-1_R | This paper |
| Sequence-based reagent | act-1_F | This paper |
| Sequence-based reagent | act-1_R | This paper |
| Sequence-based reagent | tba-1_F | This paper |
| Sequence-based reagent | tba-1_R | This paper |
| Sequence-based reagent | ubc-2_F | This paper |
| Sequence-based reagent | ubc-2_R | This paper |
| Sequence-based reagent | random primers | Invitrogen |
| Sequence-based reagent | dNTPs | Fermentas |
| plasmid | pSM | Dr. K. Mizumoto (UBC) |
| plasmid | TOPO | Invitrogen |
| Chemical compound, drug | Ampicillin | BioBasic |
| Chemical compound, drug | Carbenicillin | BioBasic |
| Chemical compound, drug | Tetracycline | BioBasic |
| Chemical compound, drug | Streptomycin | BioBasic |
| Chemical compound, drug | IPTG | Santa Cruz |
| Chemical compound, drug | SuperScript™ III Reverse Transcriptase | Invitrogen |
| Chemical compound, drug | RNAseOUT | Invitrogen |
| Chemical compound, drug | PowerTrack™ SYBR Green Master Mix | Applied Biosystems™ |
| Chemical compound, drug | Phusion Hot Start II DNA Polymerase | Thermo Scientific™ |
| Chemical compound, drug | levamisole | Sigma |
| Chemical compound, drug | Sodium azide | Thermo Scientific Chemicals |
| Chemical compound, drug | Cycloheximide | Sigma |
| Chemical compound, drug | Puromycin dihydrochloride | Sigma |
| Chemical compound, drug | N-acetyl-L-cysteine | Sigma |
| Chemical compound, drug | Ascorbic Acid (Vitamin C) | Sigma |
| Chemical compound, drug | Mitoquinone mesylate | MedChemExpress |
| Chemical compound, drug | Dihydroethidium | Sigma |
| Chemical compound, drug | CellROX Green | Invitrogen |
| Software, algorithm | ImageJ | PMID: 22930834 |
| Software, algorithm | Trimmomatic version 0.36 | PMID: 24695404 |
| Software, algorithm | Salmon version 0.9.1 | PMID: 28263959 |
| Software, algorithm | tximport | PMID: 26925227 |
| Software, algorithm | edgeR | PMID: 19910308 |
| Software, algorithm | eVITTA | PMID: 34019643 |

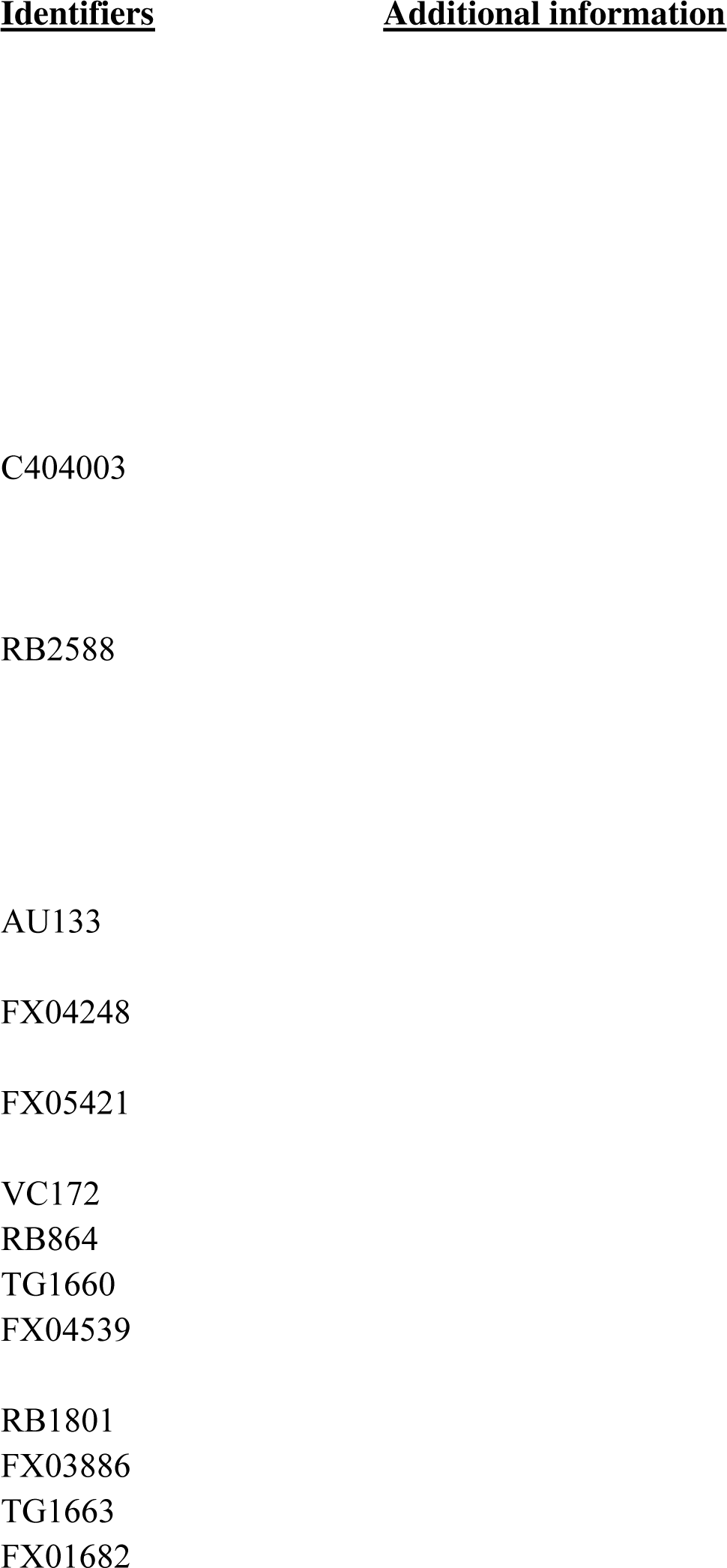
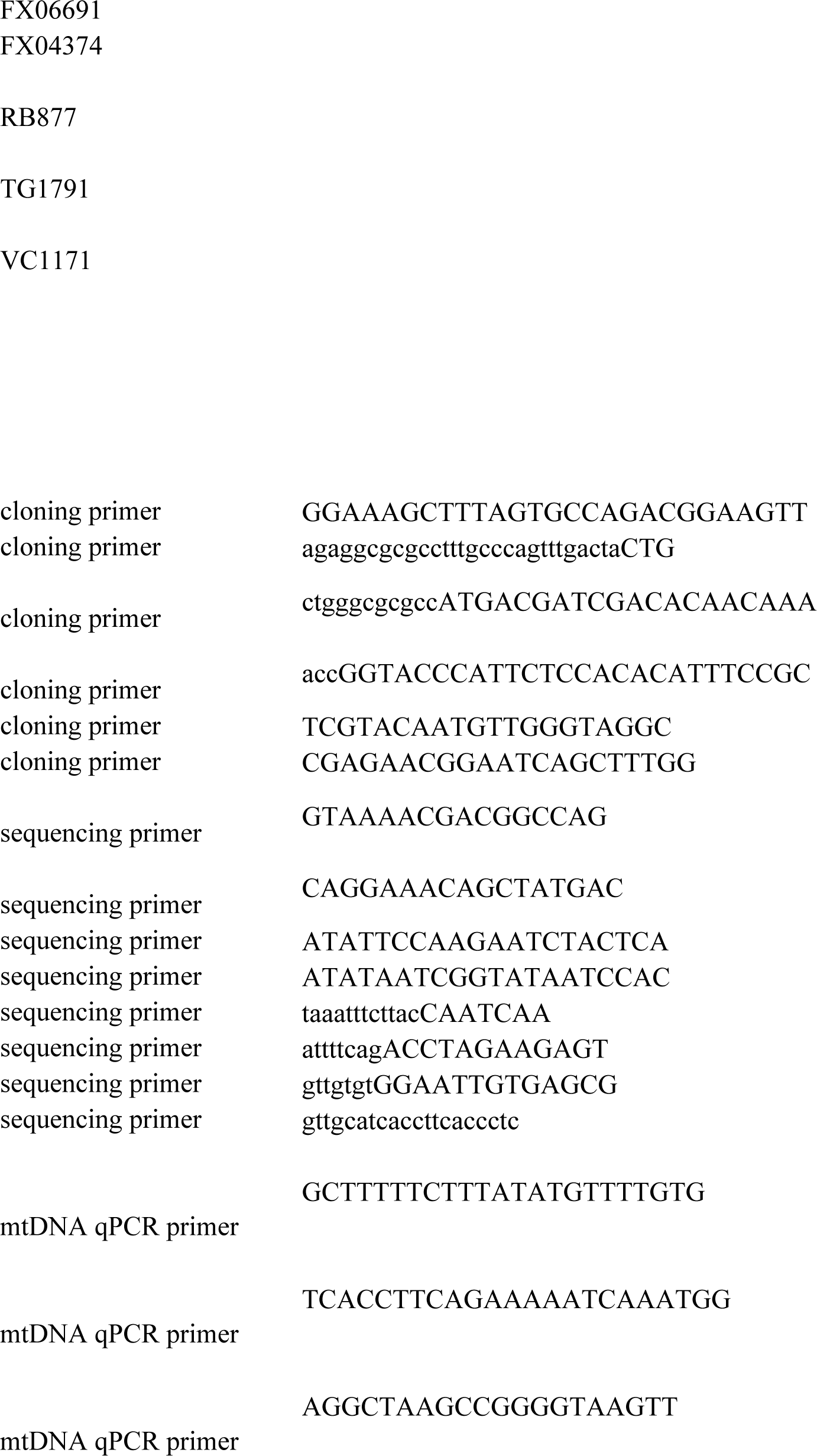
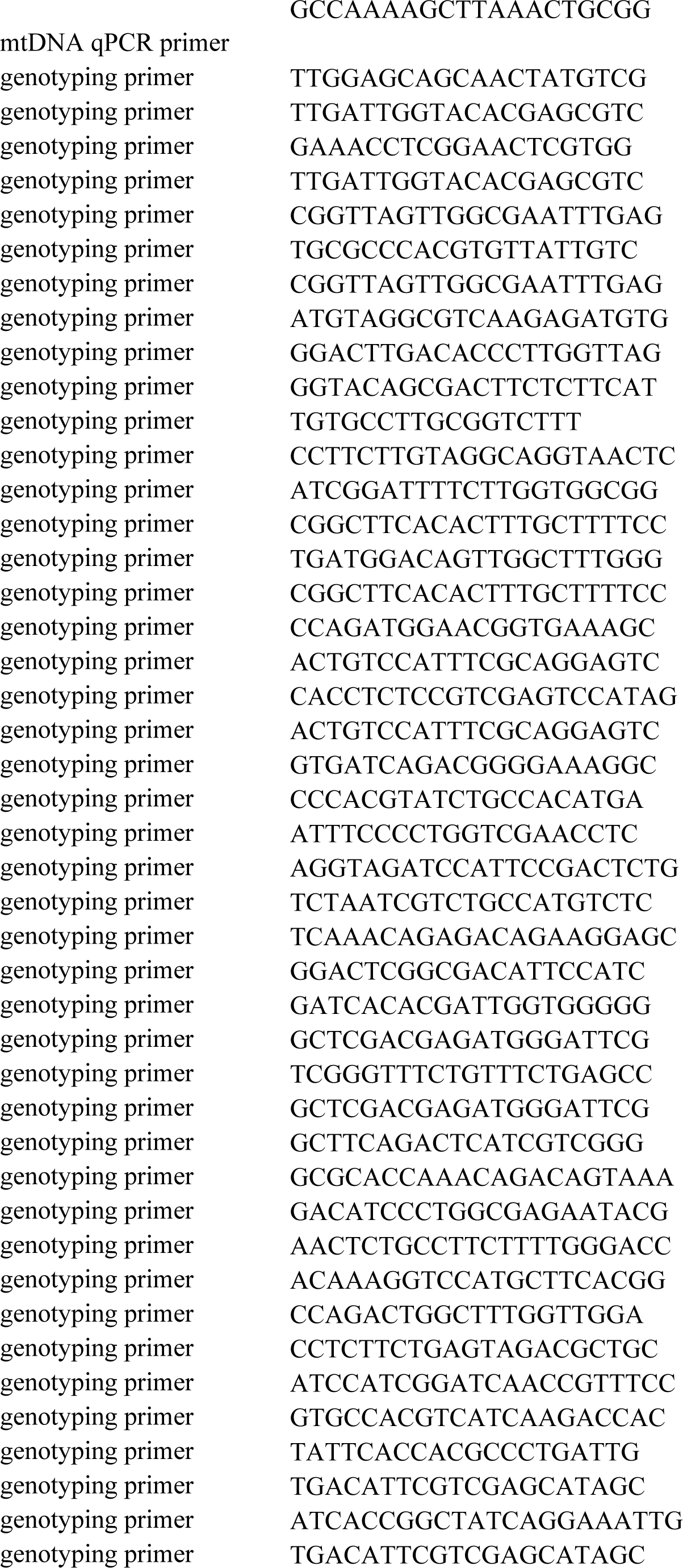
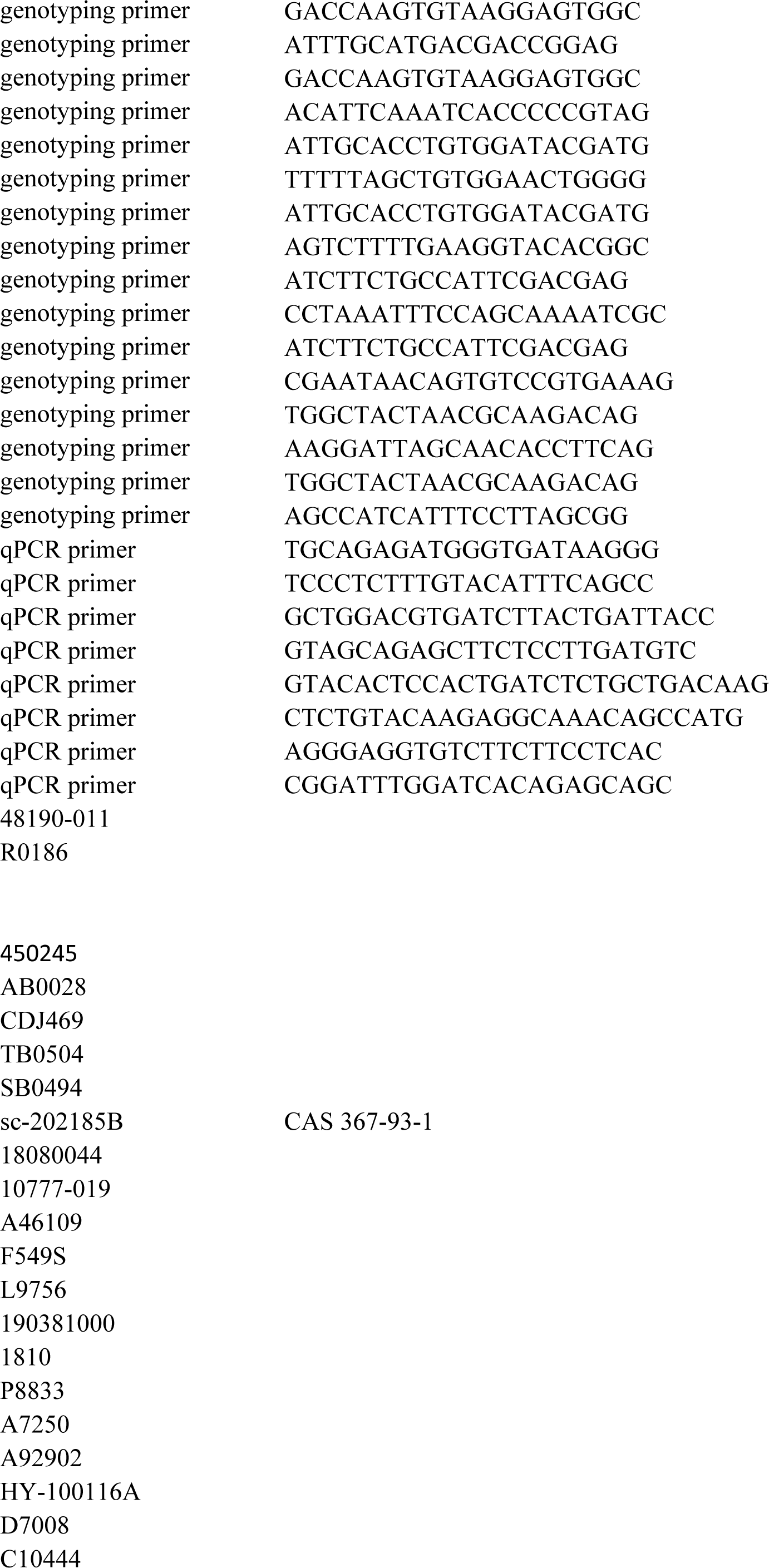
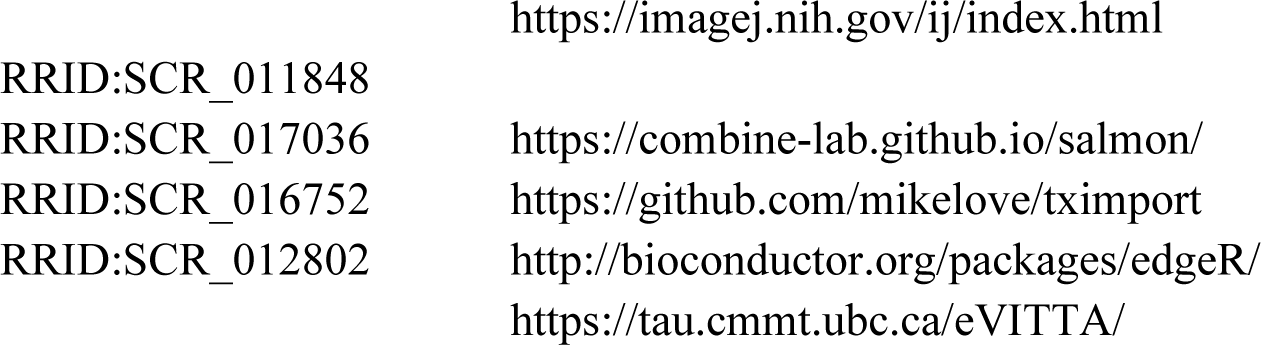

## References

1. El-Naggar AM, Sorensen PH. Translational control of aberrant stress responses as a hallmark of cancer. J Pathol. 2018;244(5):650–66.

2. Wang X, Xie J, Proud CG. Eukaryotic Elongation Factor 2 Kinase (eEF2K) in Cancer. Cancers [Internet]. 2017 Nov 27 [cited 2021 Jan 26];9(12). Available from: https://www.ncbi.nlm.nih.gov/pmc/articles/PMC5742810/

3. Dever TE, Green R. The Elongation, Termination, and Recycling Phases of Translation in Eukaryotes. Cold Spring Harb Perspect Biol. 2012 Jul;4(7):a013706.

4. Delaidelli A, Khan D, Leprivier G, Pfister SM, Taylor MD, Maris JM, et al. TB-16 exploration of eEF2K as a novel therapeutic target in medulloblastoma and neuroblastoma. Neuro-Oncol. 2016 Jun 1;18(suppl_3):iii171–iii171.

5. Leprivier G, Remke M, Rotblat B, Dubuc A, Mateo ARF, Kool M, et al. The eEF2 kinase confers resistance to nutrient deprivation by blocking translation elongation. Cell. 2013 May 23;153(5):1064–79.

6. Ryazanov AG, Shestakova EA, Natapov PG. Phosphorylation of elongation factor 2 by EF-2 kinase affects rate of translation. Nature. 1988 Jul 14;334(6178):170–3.

7. Smith PR, Loerch S, Kunder N, Stanowick AD, Lou TF, Campbell ZT. Functionally distinct roles for eEF2K in the control of ribosome availability and p-body abundance. Nat Commun. 2021 Nov 23;12(1):6789.

8. Delaidelli A, Negri GL, Jan A, Jansonius B, El-Naggar A, Lim JKM, et al. MYCN amplified neuroblastoma requires the mRNA translation regulator eEF2 kinase to adapt to nutrient deprivation. Cell Death Differ. 2017 Sep;24(9):1564–76.

9. Liu R, Proud CG. Eukaryotic elongation factor 2 kinase as a drug target in cancer, and in cardiovascular and neurodegenerative diseases. Acta Pharmacol Sin. 2016 Mar;37(3):285– 94.

10. Knight JRP, Garland G, Pöyry T, Mead E, Vlahov N, Sfakianos A, et al. Control of translation elongation in health and disease. Dis Model Mech [Internet]. 2020 Mar 1 [cited 2021 Feb 23];13(3). Available from: https://dmm.biologists.org/content/13/3/dmm043208

11. Hamurcu Z, Ashour A, Kahraman N, Ozpolat B. FOXM1 regulates expression of eukaryotic elongation factor 2 kinase and promotes proliferation, invasion and tumorgenesis of human triple negative breast cancer cells. Oncotarget. 2016 Mar 29;7(13):16619–35.

12. Zhu H, Song H, Chen G, Yang X, Liu J, Ge Y, et al. eEF2K promotes progression and radioresistance of esophageal squamous cell carcinoma. Radiother Oncol J Eur Soc Ther Radiol Oncol. 2017 Sep;124(3):439–47.

13. Xie J, Shen K, Lenchine RV, Gethings LA, Trim PJ, Snel MF, et al. Eukaryotic elongation factor 2 kinase upregulates the expression of proteins implicated in cell migration and cancer cell metastasis. Int J Cancer. 2018 May 1;142(9):1865–77.

14. Xiao M, Xie J, Wu Y, Wang G, Qi X, Liu Z, et al. The eEF2 kinase-induced STAT3 inactivation inhibits lung cancer cell proliferation by phosphorylation of PKM2. Cell Commun Signal CCS. 2020 Feb 13;18:25.

15. Lazarus MB, Levin RS, Shokat KM. Discovery of new substrates of the elongation factor-2 kinase suggests a broader role in the cellular nutrient response. Cell Signal. 2017 Jan;29:78– 83.

16. Alves V. Reactivity of vertebrate-directed phospho-eEF2 antibody against the Caenorhabditis elegans orthologue phospho-EEF-2. F1000Research. 2015 Sep 25;4:902.

17. Chu HP, Liao Y, Novak JS, Hu Z, Merkin JJ, Shymkiv Y, et al. Germline quality control: eEF2K stands guard to eliminate defective oocytes. Dev Cell. 2014 Mar 10;28(5):561–72.

18. Moore CEJ, Mikolajek H, Regufe da Mota S, Wang X, Kenney JW, Werner JM, et al. Elongation Factor 2 Kinase Is Regulated by Proline Hydroxylation and Protects Cells during Hypoxia. Mol Cell Biol. 2015 May;35(10):1788–804.

19. Láscarez-Lagunas LI, Silva-García CG, Dinkova TD, Navarro RE. LIN-35/Rb causes starvation-induced germ cell apoptosis via CED-9/Bcl2 downregulation in Caenorhabditis elegans. Mol Cell Biol. 2014 Jul;34(13):2499–516.

20. Harvald EB, Sprenger RR, Dall KB, Ejsing CS, Nielsen R, Mandrup S, et al. Multi-omics Analyses of Starvation Responses Reveal a Central Role for Lipoprotein Metabolism in Acute Starvation Survival in C. elegans. Cell Syst. 2017 Jul 26;5(1):38–52.e4.

21. Moore CEJ, Wang X, Xie J, Pickford J, Barron J, Regufe da Mota S, et al. Elongation factor 2 kinase promotes cell survival by inhibiting protein synthesis without inducing autophagy. Cell Signal. 2016 Apr;28(4):284–93.

22. Dunbar TL, Yan Z, Balla KM, Smelkinson MG, Troemel ER. C. elegans detects pathogen-induced translational inhibition to activate immune signaling. Cell Host Microbe. 2012 Apr 19;11(4):375–86.

23. Ghosh A, Singh J. Translation initiation or elongation inhibition triggers contrasting effects on Caenorhabditis elegans survival during pathogen infection [Internet]. bioRxiv; 2024 [cited 2024 Jan 27]. p. 2024.01.15.575653. Available from: https://www.biorxiv.org/content/10.1101/2024.01.15.575653v1

24. Estes KA, Dunbar TL, Powell JR, Ausubel FM, Troemel ER. bZIP transcription factor zip-2 mediates an early response to Pseudomonas aeruginosa infection in Caenorhabditis elegans. Proc Natl Acad Sci U S A. 2010 Feb 2;107(5):2153–8.

25. Schneider-Poetsch T, Ju J, Eyler DE, Dang Y, Bhat S, Merrick WC, et al. Inhibition of Eukaryotic Translation Elongation by Cycloheximide and Lactimidomycin. Nat Chem Biol. 2010 Mar;6(3):209–17.

26. Tiku V, Kew C, Mehrotra P, Ganesan R, Robinson N, Antebi A. Nucleolar fibrillarin is an evolutionarily conserved regulator of bacterial pathogen resistance. Nat Commun. 2018 Sep 6;9:3607.

27. Takauji Y, Wada T, Takeda A, Kudo I, Miki K, Fujii M, et al. Restriction of protein synthesis abolishes senescence features at cellular and organismal levels. Sci Rep. 2016 Jan 5;6(1):18722.

28. Clay KJ, Yang Y, Clark C, Petrascheck M. Proteostasis is differentially modulated by inhibition of translation initiation or elongation. Kapahi P, Manley JL, Zid BM, editors. eLife. 2023 Oct 5;12:e76465.

29. Schmidt EK, Clavarino G, Ceppi M, Pierre P. SUnSET, a nonradioactive method to monitor protein synthesis. Nat Methods. 2009 Apr;6(4):275–7.

30. Goodman CA, Hornberger TA. Measuring protein synthesis with SUnSET: a valid alternative to traditional techniques? Exerc Sport Sci Rev. 2013 Apr;41(2):107–15.

31. Aviner R. The science of puromycin: From studies of ribosome function to applications in biotechnology. Comput Struct Biotechnol J. 2020 Jan 1;18:1074–83.

32. Derisbourg MJ, Wester LE, Baddi R, Denzel MS. Mutagenesis screen uncovers lifespan extension through integrated stress response inhibition without reduced mRNA translation. Nat Commun. 2021 Mar 15;12:1678.

33. Somers HM, Fuqua JH, Bonnet FXA, Rollins JA. Quantification of tissue-specific protein translation in whole C. elegans using O-propargyl-puromycin labeling and fluorescence microscopy. Cell Rep Methods. 2022 Apr 25;2(4):100203.

34. Allavena G, Boyd C, Oo KS, Maellaro E, Zhivotovsky B, Kaminskyy VO. Suppressed translation and ULK1 degradation as potential mechanisms of autophagy limitation under prolonged starvation. Autophagy. 2016 Sep 14;12(11):2085–97.

35. Hijazi M, Casado P, Akhtar N, Alvarez-Teijeiro S, Rajeeve V, Cutillas PR. eEF2K Activity Determines Synergy to Cotreatment of Cancer Cells With PI3K and MEK Inhibitors. Mol Cell Proteomics MCP. 2022 May 2;21(6):100240.

36. Webster AK, Chitrakar R, Taylor SM, Baugh LR. Alternative somatic and germline gene-regulatory strategies during starvation-induced developmental arrest. Cell Rep. 2022 Oct 11;41(2):111473.

37. Paek J, Lo JY, Narasimhan SD, Nguyen TN, Glover-Cutter K, Robida-Stubbs S, et al. Mitochondrial SKN-1/Nrf Mediates a Conserved Starvation Response. Cell Metab. 2012 Oct 3;16(4):526–37.

38. Cui M, Cohen ML, Teng C, Han M. The Tumor Suppressor Rb Critically Regulates Starvation-Induced Stress Response in C. elegans. Curr Biol. 2013 Jun 3;23(11):975–80.

39. Cui M, Wang Y, Cavaleri J, Kelson T, Teng Y, Han M. Starvation-Induced Stress Response Is Critically Impacted by Ceramide Levels in Caenorhabditis elegans. Genetics. 2017 Feb;205(2):775–85.

40. Goh GYS, Winter JJ, Bhanshali F, Doering KRS, Lai R, Lee K, et al. NHR-49/HNF4 integrates regulation of fatty acid metabolism with a protective transcriptional response to oxidative stress and fasting. Aging Cell. 2018 Jun;17(3):e12743.

41. Derry WB, Putzke AP, Rothman JH. Caenorhabditis elegans p53: role in apoptosis, meiosis, and stress resistance. Science. 2001 Oct 19;294(5542):591–5.

42. Doering KRS, Ermakova G, Taubert S. Nuclear hormone receptor NHR-49 is an essential regulator of stress resilience and healthy aging in Caenorhabditis elegans. Front Physiol. 2023 Aug 14;14:1241591.

43. Reddy KC, Dunbar TL, Nargund AM, Haynes CM, Troemel ER. The C. elegans CCAAT-enhancer binding protein gamma is required for surveillance immunity. Cell Rep. 2016 Feb 23;14(7):1581–9.

44. McEwan DL, Kirienko NV, Ausubel FM. Host translational inhibition by Pseudomonas aeruginosa Exotoxin A Triggers an immune response in Caenorhabditis elegans. Cell Host Microbe. 2012 Apr 19;11(4):364–74.

45. Vasquez-Rifo A, Ricci EP, Ambros V. Pseudomonas aeruginosa cleaves the decoding center of Caenorhabditis elegans ribosomes. PLoS Biol. 2020 Dec 1;18(12):e3000969.

46. Hahm JH, Jeong C, Nam HG. Diet restriction-induced healthy aging is mediated through the immune signaling component ZIP-2 in Caenorhabditis elegans. Aging Cell. 2019 Oct;18(5):e12982.

47. Jones RG, Plas DR, Kubek S, Buzzai M, Mu J, Xu Y, et al. AMP-Activated Protein Kinase Induces a p53-Dependent Metabolic Checkpoint. Mol Cell. 2005 Apr 29;18(3):283–93.

48. Lacroix M, Riscal R, Arena G, Linares LK, Le Cam L. Metabolic functions of the tumor suppressor p53: Implications in normal physiology, metabolic disorders, and cancer. Mol Metab. 2020 Mar 1;33:2–22.

49. Lim JK, Samiei A, Carnie CJ, Brinkman V, Radiloff D, Cran J, et al. The eEF2 kinase coordinates the DNA damage response to cisplatin by supporting p53 activation [Internet]. bioRxiv; 2023 [cited 2024 Jan 27]. p. 2023.03.28.534603. Available from: https://www.biorxiv.org/content/10.1101/2023.03.28.534603v1

50. Hofmann ER, Milstein S, Boulton SJ, Ye M, Hofmann JJ, Stergiou L, et al. Caenorhabditis elegans HUS-1 Is a DNA Damage Checkpoint Protein Required for Genome Stability and EGL-1-Mediated Apoptosis. Curr Biol. 2002 Nov 19;12(22):1908–18.

51. The C. elegans Deletion Mutant Consortium. Large-Scale Screening for Targeted Knockouts in the Caenorhabditis elegans Genome. G3 GenesGenomesGenetics. 2012 Nov 1;2(11):1415–25.

52. Van Gilst MR, Hadjivassiliou H, Yamamoto KR. A Caenorhabditis elegans nutrient response system partially dependent on nuclear receptor NHR-49. Proc Natl Acad Sci U S A. 2005 Sep 20;102(38):13496–501.

53. Baugh LR, Hu PJ. Starvation Responses Throughout the Caenorhabditis elegans Life Cycle. Genetics. 2020 Dec;216(4):837–78.

54. Kumar N, Raja S, Van Houten B. The involvement of nucleotide excision repair proteins in the removal of oxidative DNA damage. Nucleic Acids Res. 2020 Nov 18;48(20):11227–43.

55. Lans H, Vermeulen W. Nucleotide Excision Repair in Caenorhabditis elegans. Mol Biol Int. 2011;2011:542795.

56. Chatterjee N, Walker GC. Mechanisms of DNA damage, repair and mutagenesis. Environ Mol Mutagen. 2017 Jun;58(5):235–63.

57. Volkova NV, Meier B, González-Huici V, Bertolini S, Gonzalez S, Vöhringer H, et al. Mutational signatures are jointly shaped by DNA damage and repair. Nat Commun. 2020 May 1;11(1):2169.

58. Sabatella M, Thijssen KL, Davó-Martínez C, Vermeulen W, Lans H. Tissue-Specific DNA Repair Activity of ERCC-1/XPF-1. Cell Rep [Internet]. 2021 Jan 12 [cited 2022 Jan 6];34(2). Available from: https://www.cell.com/cell-reports/abstract/S2211-1247(20)31597-7

59. Astin JW, O’Neil NJ, Kuwabara PE. Nucleotide excision repair and the degradation of RNA pol II by the Caenorhabditis elegans XPA and Rsp5 orthologues, RAD-3 and WWP-1. DNA Repair. 2008 Feb 1;7(2):267–80.

60. Elsakrmy N, Zhang-Akiyama QM, Ramotar D. The Base Excision Repair Pathway in the Nematode Caenorhabditis elegans. Front Cell Dev Biol. 2020 Dec 3;8:598860.

61. Roux AE, Langhans K, Huynh W, Kenyon C. Reversible Age-Related Phenotypes Induced during Larval Quiescence in C. elegans. Cell Metab. 2016 Jun 14;23(6):1113–26.

62. Tao J, Wu QY, Ma YC, Chen YL, Zou CG. Antioxidant response is a protective mechanism against nutrient deprivation in C. elegans. Sci Rep. 2017 Feb 23;7:43547.

63. Hibshman JD, Leuthner TC, Shoben C, Mello DF, Sherwood DR, Meyer JN, et al. Nonselective autophagy reduces mitochondrial content during starvation in Caenorhabditis elegans. Am J Physiol Cell Physiol. 2018 Dec 1;315(6):C781–92.

64. Edifizi D, Nolte H, Babu V, Castells-Roca L, Mueller MM, Brodesser S, et al. Multilayered Reprogramming in Response to Persistent DNA Damage in C. elegans. Cell Rep. 2017 Aug 29;20(9):2026–43.

65. Mueller MM, Castells-Roca L, Babu V, Ermolaeva MA, Müller RU, Frommolt P, et al. DAF-16/FOXO and EGL-27/GATA promote developmental growth in response to persistent somatic DNA damage. Nat Cell Biol. 2014 Dec;16(12):1168–79.

66. Murphy MP. How mitochondria produce reactive oxygen species. Biochem J. 2009 Jan 1;417(Pt 1):1–13.

67. Macedo F, Martins GL, Luévano-Martínez LA, Viana GM, Riske KA, Inague A, et al. Lipase-like 5 enzyme controls mitochondrial activity in response to starvation in Caenorhabditis elegans. Biochim Biophys Acta BBA - Mol Cell Biol Lipids. 2020 Feb 1;1865(2):158539.

68. Kelso GF, Porteous CM, Coulter CV, Hughes G, Porteous WK, Ledgerwood EC, et al. Selective Targeting of a Redox-active Ubiquinone to Mitochondria within Cells: ANTIOXIDANT AND ANTIAPOPTOTIC PROPERTIES*. J Biol Chem. 2001 Feb 16;276(7):4588–96.

69. Ng LF, Gruber J, Cheah IK, Goo CK, Cheong WF, Shui G, et al. The mitochondria-targeted antioxidant MitoQ extends lifespan and improves healthspan of a transgenic Caenorhabditis elegans model of Alzheimer disease. Free Radic Biol Med. 2014 Jun 1;71:390–401.

70. Tsui KH, Li CJ. Mitoquinone shifts energy metabolism to reduce ROS-induced oxeiptosis in female granulosa cells and mouse oocytes. Aging. 2023 Jan 9;15(1):246–60.

71. Coppa A, Guha S, Fourcade S, Parameswaran J, Ruiz M, Moser AB, et al. The peroxisomal fatty acid transporter ABCD1/PMP-4 is required in the C. elegans hypodermis for axonal maintenance: A worm model for adrenoleukodystrophy. Free Radic Biol Med. 2020 May 20;152:797–809.

72. Hoffman S, Martin D, Meléndez A, Bargonetti J. C. elegans CEP-1/p53 and BEC-1 Are Involved in DNA Repair. PLOS ONE. 2014 Feb 20;9(2):e88828.

73. Liao Y, Chu HP, Hu Z, Merkin JJ, Chen J, Liu Z, et al. Paradoxical Roles of Elongation Factor-2 Kinase in Stem Cell Survival. J Biol Chem. 2016 Sep 9;291(37):19545–57.

74. Derry WB, Bierings R, van Iersel M, Satkunendran T, Reinke V, Rothman JH. Regulation of developmental rate and germ cell proliferation in Caenorhabditis elegans by the p53 gene network. Cell Death Differ. 2007 Apr;14(4):662–70.

75. Kuo CT, You GT, Jian YJ, Chen TS, Siao YC, Hsu AL, et al. AMPK-mediated formation of stress granules is required for dietary restriction-induced longevity in Caenorhabditis elegans. Aging Cell. 2020;19(6):e13157.

76. Brenner S. The Genetics of CAENORHABDITIS ELEGANS. Genetics. 1974 May;77(1):71–94.

77. Doering KR, Cheng X, Milburn L, Ratnappan R, Ghazi A, Miller DL, et al. Nuclear hormone receptor NHR-49 acts in parallel with HIF-1 to promote hypoxia adaptation in Caenorhabditis elegans. Portman D, Sengupta P, editors. eLife. 2022 Mar 14;11:e67911.

78. Hendi A, Mizumoto K. GFPnovo2, a brighter GFP variant for in vivo labeling in C. elegans. MicroPublication Biol [Internet]. 2018 Sep 4 [cited 2024 Jan 5]; Available from: https://www.micropublication.org/journals/biology/49yb-7k39

79. Lee BH, Ashrafi K. A TRPV channel modulates C. elegans neurosecretion, larval starvation survival, and adult lifespan. PLoS Genet. 2008 Oct;4(10):e1000213.

80. Sommer I, Schwebel H, Adamo V, Bonnabry P, Bouchoud L, Sadeghipour F. Stability of N-Acetylcysteine (NAC) in Standardized Pediatric Parenteral Nutrition and Evaluation of N,N-Diacetylcystine (DAC) Formation. Nutrients. 2020 Jun 21;12(6):1849.

81. Yin X, Chen K, Cheng H, Chen X, Feng S, Song Y, et al. Chemical Stability of Ascorbic Acid Integrated into Commercial Products: A Review on Bioactivity and Delivery Technology. Antioxidants. 2022 Jan 13;11(1):153.

82. van der Woude M, Lans H. C. elegans survival assays to discern global and transcription-coupled nucleotide excision repair. STAR Protoc. 2021 Jun 8;2(2):100586.

83. Misare KR, Ampolini EA, Gonzalez HC, Sullivan KA, Li X, Miller C, et al. The consequences of tetraploidy on Caenorhabditis elegans physiology and sensitivity to chemotherapeutics. Sci Rep. 2023 Oct 23;13:18125.

84. Goh GYS, Martelli KL, Parhar KS, Kwong AWL, Wong MA, Mah A, et al. The conserved Mediator subunit MDT-15 is required for oxidative stress responses in *Caenorhabditis elegans*. Aging Cell. 2014 Feb;13(1):70–9.

85. Bolger AM, Lohse M, Usadel B. Trimmomatic: a flexible trimmer for Illumina sequence data. Bioinformatics. 2014 Aug 1;30(15):2114–20.

86. Patro R, Duggal G, Love MI, Irizarry RA, Kingsford C. Salmon: fast and bias-aware quantification of transcript expression using dual-phase inference. Nat Methods. 2017 Apr;14(4):417.

87. Soneson C, Love MI, Robinson MD. Differential analyses for RNA-seq: transcript-level estimates improve gene-level inferences. F1000Research [Internet]. 2015 [cited 2020 Dec 2];4. Available from: http://www.ncbi.nlm.nih.gov/pmc/articles/PMC4712774/

88. Robinson MD, McCarthy DJ, Smyth GK. edgeR: a Bioconductor package for differential expression analysis of digital gene expression data. Bioinformatics. 2010 Jan 1;26(1):139– 40.

89. Cheng X, Yan J, Liu Y, Wang J, Taubert S. eVITTA: a web-based visualization and inference toolbox for transcriptome analysis. Nucleic Acids Res. 2021 Jul 2;49(W1):W207– 15.

90. Goh GYS, Beigi A, Yan J, Doering KRS, Taubert S. Mediator subunit MDT-15 promotes expression of propionic acid breakdown genes to prevent embryonic lethality in Caenorhabditis elegans. G3 GenesGenomesGenetics. 2023 Jun 1;13(6):jkad087.

91. Shpilka T, Du Y, Yang Q, Melber A, Uma Naresh N, Lavelle J, et al. UPRmt scales mitochondrial network expansion with protein synthesis via mitochondrial import in Caenorhabditis elegans. Nat Commun. 2021 Jan 20;12:479.

