## Supplementary Figures for "Eukaryotic Elongation Factor 2 Kinase EFK-1/eEF2K promotes starvation resistance by preventing oxidative damage in *C. elegans*"

### Supplementary information

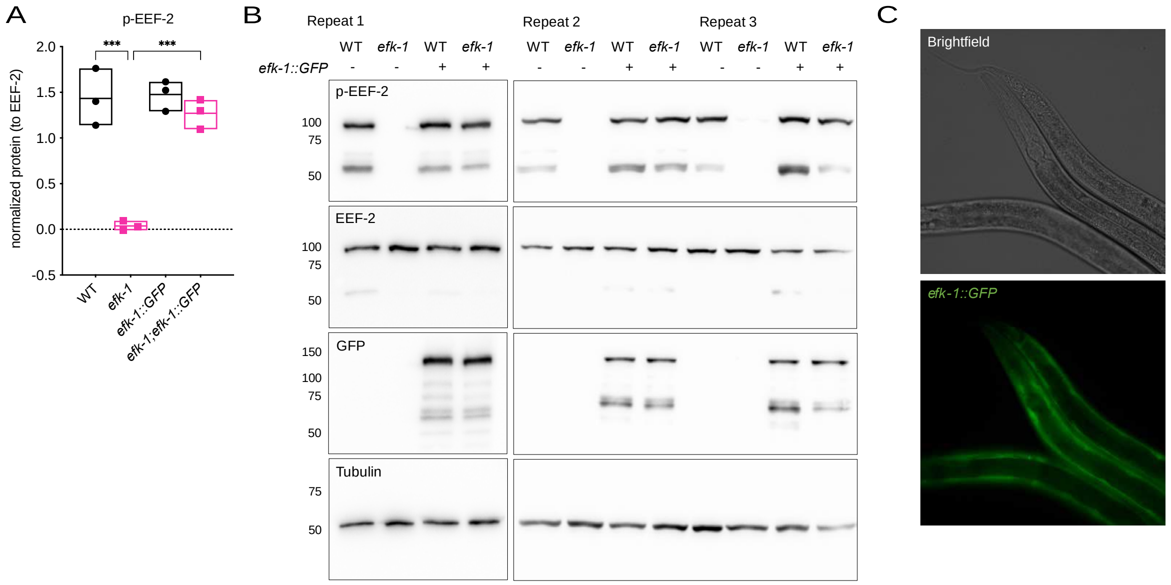


**Figure S1. (A)** Quantification of p-EEF-2 in Figure 1B. Background-corrected signal is normalized to EEF-2 control. N=3; ***p<0.001 (one-way ANOVA with Tukey's multiple comparisons test). **(B)** Full WB membranes corresponding to Figure 1B. **(C)** Brightfield and fluorescence micrographs of *efk-1::GFP* expressing transgenic worms at L4 stage. WT, wild-type. See Source data for **(A)**.

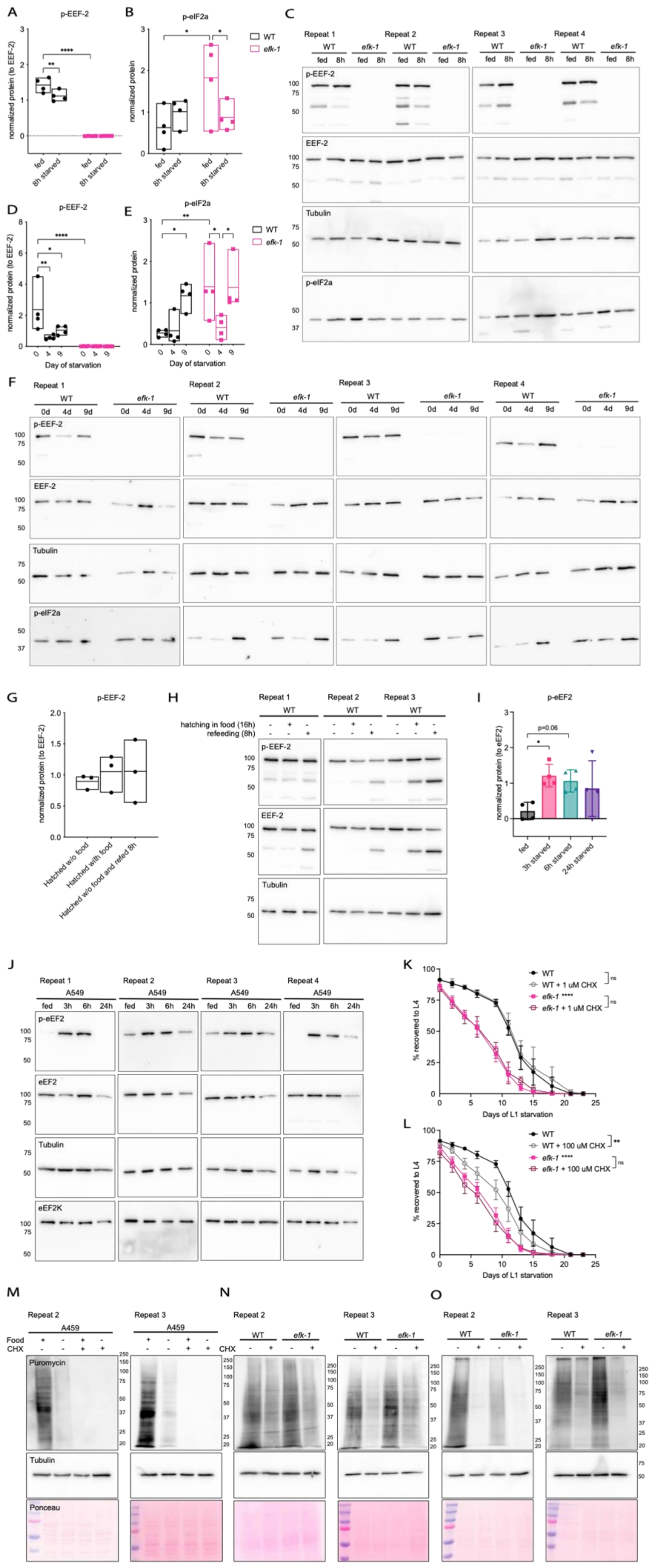
A
p-EEF-2
normalized protein (to EEF-2)
fed
8h starved
B
p-eIF2a
normalized protein
WT
efk-1
C
Repeat 1
Repeat 2
Repeat 3
Repeat 4
p-EEF-2
EEF-2
Tubulin
p-eIF2a
D
p-EEF-2
Day of starvation
E
p-eIF2a
F
G
Hatched w/o food
Hatched with food
Hatched w/o food and refed 8h
H
hatching in food (16h)
refeeding (8h)
I
p-eEF2
normalized protein (to eEF2)
p=0.06
fed
3h starved
6h starved
24h starved
J
A549
fed
3h
6h
24h
p-eEF2
eEF2
Tubulin
eEF2K
K
WT
WT + 1 uM CHX
efk-1 ****
efk-1 + 1 uM CHX
ns
% recovered to L4
Days of L1 starvation
L
WT + 100 uM CHX
efk-1 + 100 uM CHX
**
M
A459
Food
CHX
Puromycin
Ponceau
N
O

**Figure S2. (A-B)** Quantification of **(A)** p-EEF-2 and **(B)** p-eIF2$\alpha$ in Figure 2A, normalized to EEF-2 and tubulin, respectively. N=4; *$p<0.05$, **$p<0.01$, ****$p<0.0001$ (two-way ANOVA with uncorrected Fisher's LSD). **(C)** Full WB membranes corresponding to Figure 2A. **(D-E)** Quantification of **(D)** p-EEF-2 and **(E)** p-eIF2$\alpha$ in Figure 2B, normalized to EEF-2 and tubulin, respectively. N=4; *$p<0.05$, **$p<0.01$, ****$p<0.0001$ (two-way ANOVA with Tukey’s multiple comparisons test). **(F)** Full WB membranes corresponding to Figure 2B. **(G)** Quantification of p-EEF-2 in Figure 2C, normalized to EEF-2. N=3; no significant comparisons (one-way ANOVA with Dunnett’s multiple comparisons test). **(H)** Full WB membranes corresponding to Figure 2C. **(I)** Quantification of p-eEF2 in Figure 2D, normalized to eEF2. N=4, error bars represent SD; *$p<0.05$ (one-way ANOVA with Dunnett’s multiple comparisons test). **(J)** Full WB membranes corresponding to Figure 2D. **(K-L)** The graphs show L1 starvation survival of WT and *efk-1* mutants with or without supplementation of CHX at **(K)** 1 uM and **(L)** 100 uM. N=4, error bars represent SD; ****$p<0.0001$ percent L4 vs. WT animals (AUC compared using one-way ANOVA with Tukey’s multiple comparisons test). Controls shown are of the same experiment as Figure 2F. **(M-O)** Full WB membranes and Ponceau S staining for additional 2 biological repeats of **(M)** fed and starved A549 cells (Figure 2G), **(N)** 8-hour starved L4 worms (Figure 2H), and **(O)** 6-day starved L1 worms (Figure 2I), respectively. N=3 total shown for each experiment. WT, wild-type; ns, not significant; AUC, area under the curve; CHX, cycloheximide. See Source data for **(A-B, D-E, G, I, K-L)**.

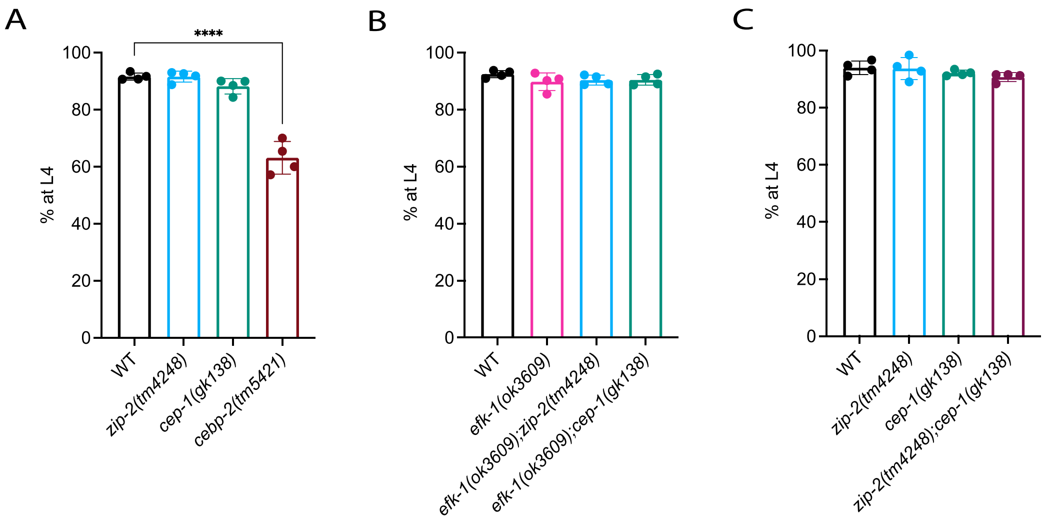


**Figure S3. (A)** The graph shows the growth rates of WT, *zip-2* and *cep-1* and *cebp-2* mutants in unstressed conditions. N=4, error bars represent SD; ****p<0.0001 percent L4 vs. WT (one-way ANOVA with Dunnett's multiple comparisons test). **(B-C)** The graph shows the growth rates of **(B)** *efk-1*, *efk-1;zip-2* and *efk-1;cep-1* mutants, and **(C)** *zip-2*, *cep-1* and *zip-2;cep-1* mutants in unstressed conditions. N=4, error bars represent SD; no significant comparisons vs. WT (one-way ANOVA with Dunnett's multiple comparisons test). WT, wild-type. See Source data for **(A-C)**.

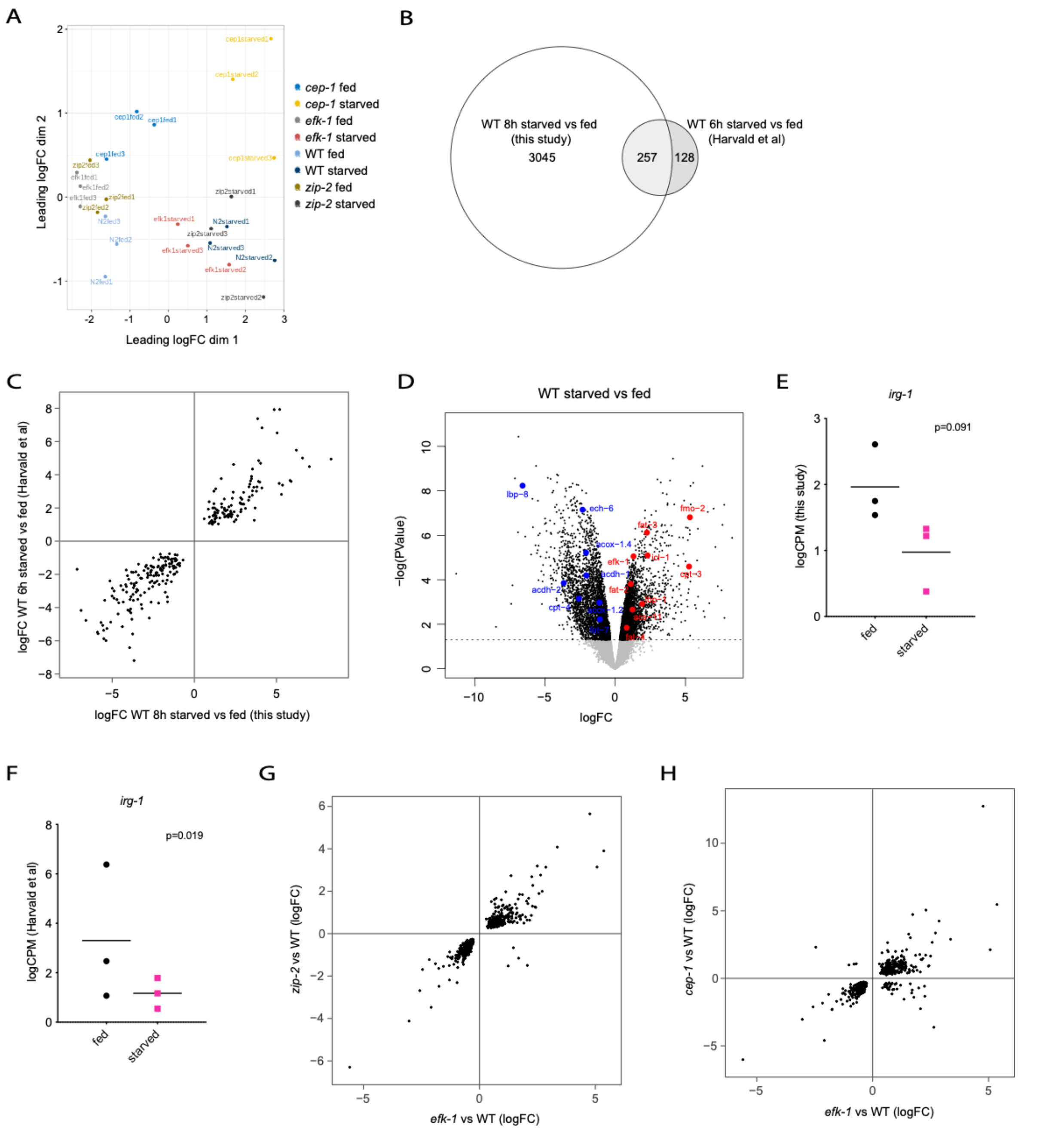


**Figure S4. (A)** The MDS plot shows the spread of expression patterns in RNA-seq samples of fed and starved WT and *efk-1*, *zip-2* and *cep-1* mutant worms. **(B)** The Venn diagram shows overlap of DEGs ($p<0.005$, $FDR<0.05$) in starved WT in this study (L4 8-hour starved vs fed) and Harvald et al. (L4 6-hour starved vs fed) (20). p=7.62e-128. **(C)** The scatter plot shows correlation of DEGs ($p<0.005$, $FDR<0.05$) in this study and Harvald et al. (20). n=257, $r^2=0.87$, p=2.2e-16. **(D)** The Volcano plot of differentially expressed genes in WT starved vs fed in our study. x=logFC, y=-log(PValue). Red, upregulated genes; blue; downregulated genes (p<0.05). **(E-F)** The logCPM plots show *irg-1* expression in fed and starved worms in **(E)** this study and **(F)** Harvald et al. (20). P-value from corresponding DEG analysis are shown. **(G-H)** The scatter plots show correlation of DEGs in **(G)** *zip-2* vs *efk-1* and **(H)** *cep-1* vs *efk-1*. x= ES (x), y= ES (y). $r^2$=0.84, p<2.2e-16 for *zip-2* vs *efk-1*; $r^2$=0.57, p<2.2e-16 for *cep-1* vs *efk-1*. MDS, Multi-dimensional scaling; DEGs, differentially expressed genes; FDR, false discovery rate; CPM, counts per million. See Source data for **(E-F)**.

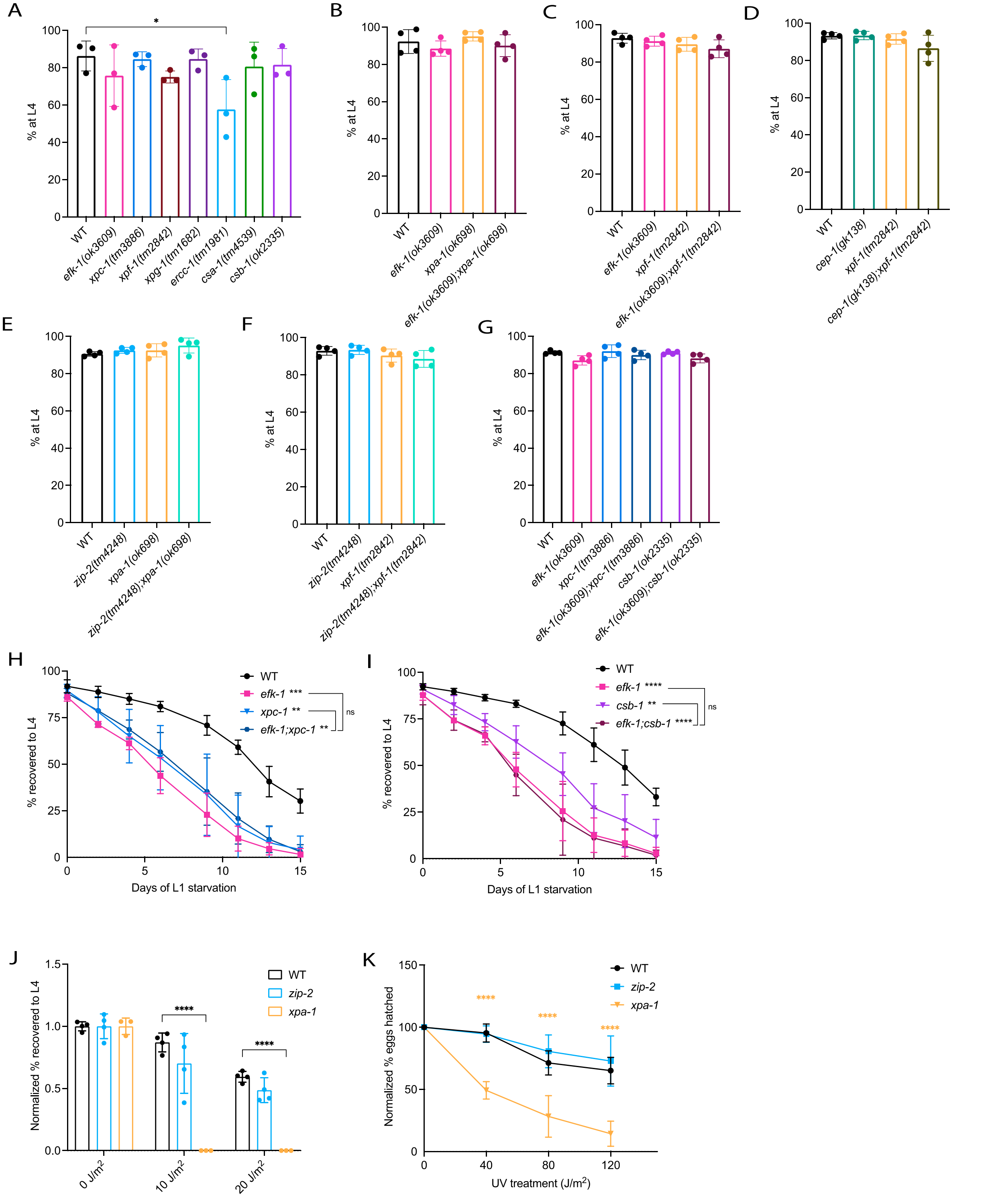


**Figure S5. (A)** The graph shows the growth rates of *xpc-1*, *xpf-1*, *xpg-1*, *csa-1*, *csb-1* and *ercc-1* worms in unstressed conditions. N=3, error bars represent SD; *p<0.05 percent L4 vs.

WT (one-way ANOVA with Dunnett's multiple comparisons test). **(B-G)** The graphs show the growth rates of double mutants *efk-1;xpa-1*, *efk-1;xpf-1*, *cep-1;xpf-1*, *zip-2;xpa-1*, *zip-2;xpf-1*, *efk-1;xpc-1 efk-1;csb-1* and respective single mutants under normal conditions. N=4, error bars represent SD; no significant comparisons (one-way ANOVA with Dunnett's multiple comparisons test). **(H-I)** The graphs show L1 starvation survival of **(H)** *efk-1;xpc-1* and **(I)** *efk-1;csb-1* double mutants alongside respective single mutants and WT control. N=4, error bars represent SD; **p<0.01, ***p<0.001, ****p<0.0001 percent L4 vs. WT (AUC compared using one-way ANOVA with Tukey's multiple comparisons test). **(J-K)** The graphs show the larval and embryonic UV-C resistance of *zip-2* mutants, as measured by **(J)** recovery to L4 stage after UVC irradiation (0-20 $J/m^2$) at L1, and **(K)** percentages of viable embryos (24h after egglay) after parents were subjected to UVC irradiation (0-120 $J/m^2$) at young adult stage. Data are normalized to no UV control. WT and *xpa-1* controls shown are of the same experiment as Figure 5I-L. N=4, error bars represent SD; ****p<0.0001 (two-way ANOVA with Dunnett's multiple comparisons test). WT, wild-type; ns, not significant; AUC, area under the curve. See Source data for **(A-K)**.

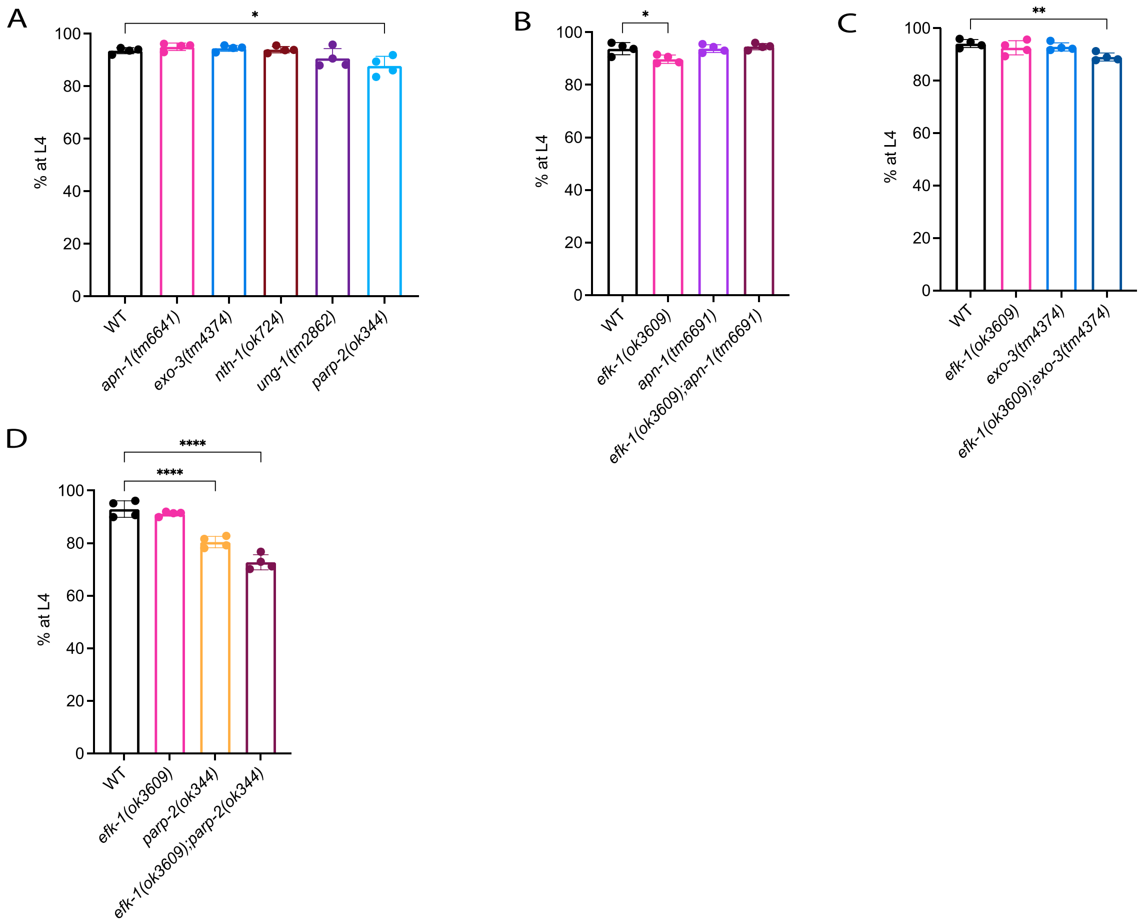


**Figure S6. (A)** The graphs show the growth rates of WT, *apn-1*, *exo-3*, *nth-1*, *ung-1* and *parp-2* mutants in unstressed conditions. N=4, error bars represent SD; *p<0.05 (one-way ANOVA with Dunnett's multiple comparisons test). **(B-D)** The graphs show the growth rates of double mutants *efk-1;apn-1*, *efk-1;exo-3*, *efk-1;parp-2*, respective single mutants and WT control. N=4, error bars represent SD; *p<0.05, **p<0.01, ****p<0.0001 (one-way ANOVA with Dunnett's multiple comparisons test). WT, wild-type. See Source data for **(A-D)**.

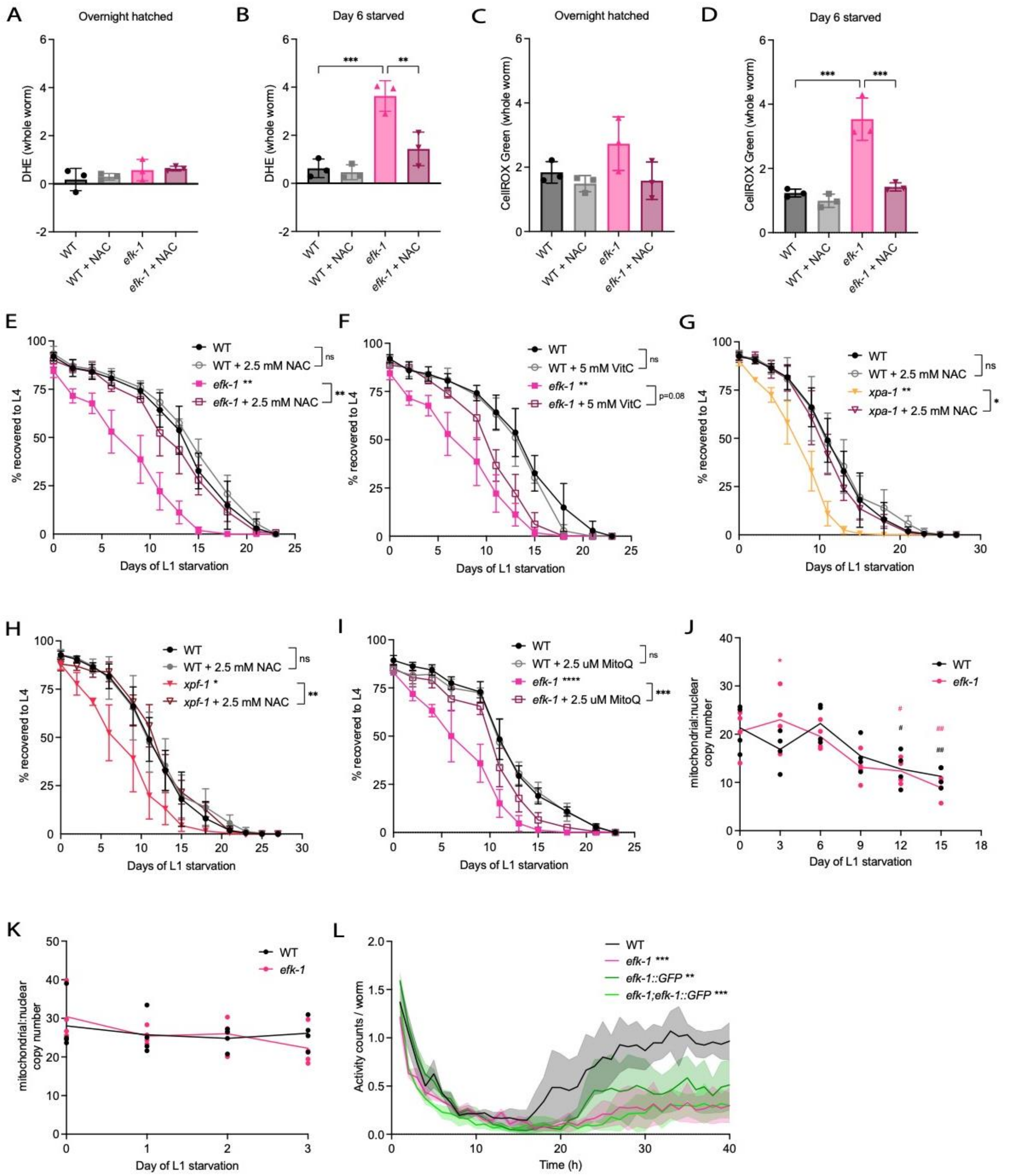


**Figure S7. (A-B)** Quantification of DHE signal from Figure 7A in **(A)** overnight hatched, and **(B)** day 6 starved samples. **(C-D)** Quantification of CellROX Green signal from Figure 7B in **(C)** overnight hatched, and **(D)** day 6 starved samples. N=3 (average of 30~40 worms per repeat), error bars represent SD; **p<0.01, ***p<0.001 (one-way ANOVA with Tukey's multiple comparisons test). **(E-F)** The graphs show L1 starvation survival of WT and *efk-1* mutants with or without supplementation of antioxidants **(E)** NAC and **(F)** VitC at 2.5 and 5 mM, respectively. WT and *efk-1* controls shown are of the same experiments as Figure 7C-D. N=3, error bars represent SD; **p<0.01 (AUC compared using one-way ANOVA with Tukey's multiple comparisons test). **(G-H)** The graphs show L1 starvation survival of WT and **(G)** *xpa-1* or **(H)** *xpf-1* mutants with or without supplementation of 2.5 mM NAC. WT and WT+NAC controls shown are of the same experiments as Figure 7E-F. N=4, error bars represent SD; *p<0.05, **p<0.01 (AUC compared using one-way ANOVA with Tukey's multiple comparisons test). **(I)** The graph shows the L1 starvation survival of WT and *efk-1* mutants with or without supplementation of mitochondrial antioxidant MitoQ at 2.5 uM. N=4, error bars represent SD; ***p<0.001, ****p<0.0001 (AUC compared using one-way ANOVA with Tukey's multiple comparisons test). **(J-K)** The graphs show mitochondrial DNA copy number of WT and *efk-1* mutants in **(J)** prolonged (up to 15 days) and **(K)** early (0-3 days) L1 starvation. N=4, *p<0.05 vs. WT, $^{\#}$p<0.05, $^{\#\#}$p<0.01 vs. day 0 (newly hatched) samples (two-way ANOVA with Tukey's multiple comparisons test). **(L)** The graph shows activity counts per worm (y-axis) of WT and *efk-1* mutants with or without the rescue construct *efk-1::GFP* in L1 starvation (0~40h, x-axis). N=3, shaded area represents SEM; **p<0.01, ***p<0.001 (one-way ANOVA with Tukey's multiple comparisons test). WT, wild-type; NAC, N-acetyl-L-cysteine; VitC, Vitamin C (ascorbic acid); ns, not significant; AUC, area under the curve; MitoQ, mitoquinone. See Source data for **(A-L)**.
